# Convergent and divergent structural and functional brain abnormalities associated with developmental dyslexia: a cross-linguistic meta-analysis of neuroimaging studies

**DOI:** 10.1101/2021.05.10.443380

**Authors:** Xiaohui Yan, Ke Jiang, Hui Li, Ziyi Wang, Kyle Perkins, Fan Cao

**Affiliations:** 1 Department of Psychology, Sun Yat-Sen University; 2 Anyang Preschool Education College; 3 Foreign Language Department, Jining University; 4 Florida International University (Retired Professor)

**Keywords:** dyslexia, fMRI/PET, VBM, meta-analysis, multimodal, alphabetic language, morpho-syllabic language

## Abstract

Brain abnormalities in the reading network have been repeatedly reported in individuals with developmental dyslexia (DD); however, it is still not totally understood where and why the structural and functional abnormalities are consistent/inconsistent across languages. In the current multimodal meta-analysis, we found convergent structural and functional alterations in the left superior temporal gyrus across languages, suggesting a neural signature of DD. We found greater reduction in grey matter volume and brain activation in the left inferior frontal gyrus in morpho-syllabic languages (e.g. Chinese) than in alphabetic languages, and greater reduction in brain activation in the left middle temporal gyrus and fusiform gyrus in alphabetic languages than in morpho-syllabic languages. These language differences are explained as consequences of being DD while learning a specific language. In addition, we also found brain regions that showed increased grey matter volume and brain activation, presumably suggesting compensations and brain regions that showed inconsistent alterations in brain structure and function. Our study provides important insights about the etiology of DD from a cross-linguistic perspective with considerations of consistency/inconsistency between structural and functional alterations.

## Introduction

Individuals with developmental dyslexia (DD) encounter difficulty in learning to read even with normal intelligence and adequate educational guidance (Peterson & Pennington, 2012). DD affects a large number of individuals across writing systems, and the prevalence is about 5-10% in alphabetic writing systems (e.g., English and German) (Döhla & Heim, 2016; Katusic et al., 2001; Shaywitz, 1996) and about 4-7% in morpho-syllabic writing systems (e.g., Chinese and Japanese Kanji) (Sun et al., 2013; Uno et al., 2009; Zhao et al., 2016). A phonological deficit in DD has been documented across languages (Gu & Bi, 2020; Snowling & Melby-Lervag, 2016). Individuals with DD show deficient phonological ability including phonological representation, manipulation and retrieval even when compared to reading-level controls (Melby-Lervag et al., 2012; Parrila et al., 2020). However, the common phonological deficit may manifest differently in reading behavior depending on the specific requirements of the writing system. For example, phonological deficit in English is associated with lower accuracy in phonological decoding (Landerl et al., 1997; Ziegler et al., 2003), and it is associated with slower reading speed in transparent orthographies with relatively intact accuracy in phonological decoding (Wimmer & Schurz, 2010). In Chinese, phonological deficit is associated with a higher rate of semantic errors during character reading (Shu et al., 2005), because children with DD over-rely on the semantic cue in the character during reading due to the inability to use the phonological cue. According to research, 80% of Chinese characters have a semantic radical and a phonetic radical providing semantic cues and phonological cues of the character, respectively (Honorof & Feldman, 2006).

At the neurological level, individuals with DD are associated with dysfunction in the left reading network (Richlan, 2012), including the occipitotemporal cortex (OT), the temporoparietal cortex (TP) and the inferior frontal cortex. The left TP area, including the posterior superior temporal gyrus (STG) and inferior parietal lobule (IPL), is a key area related to phonological representation and phonological conversion (Petersen & Fiez, 1993; Richlan, 2012). This region tends to show reduced brain activation in alphabetic languages as demonstrated in a cross-linguistic study of English, Italian and French (Paulesu et al., 2001) and several meta-analysis studies in alphabetic languages (Maisog et al., 2008; Martin et al., 2016; Paulesu et al., 2014; Richlan et al., 2009, 2011). The left OT area, including the middle occipital gyrus (MOG), inferior temporal gyrus (ITG) and fusiform gyrus, has been consistently found to show reduced activation in individuals with DD across morpho-syllabic and alphabetic languages (Bolger et al., 2008; Cao et al., 2020; Centanni et al., 2019; Chyl et al., 2018; Paz-Alonso et al., 2018). This region is associated with visuo-orthographic processing during reading (Glezer et al., 2016., 2019). The left inferior frontal gyrus (IFG) has been known to be involved in phonological and semantic retrieval, lexical selection and integration (Booth et al., 2007a, 2007b; Costafreda et al., 2006; Szatkowska et al., 2000). However, the nature of dysfunction in the left IFG in individuals with DD remains controversial. Although reduced activation in the left IFG was confirmed by many fMRI studies and meta-analysis studies (Booth et al., 2007a; Cao et al., 2006; Richlan et al., 2010; Wimmer et al., 2010), increased activation in the left IFG was also reported in many fMRI studies (Grunling et al., 2004; Kronbichler et al., 2006; Waldie et al., 2013; Wimmer et al., 2010). The inconsistent results may be related to task difficulty (Waldie et al., 2013; Wimmer et al., 2010), orthographic transparency (Martin et al., 2016) and age of participants (Chyl et al., 2018).

From a cross-linguistic perspective, quite a few studies have found language-universal deficits associated with DD. Paulesu et al. (2001) found that readers with DD in English, Italian and French showed similar brain abnormality during an explicit word reading task and an implicit reading task, namely, reduced brain activation at the left middle/superior temporal gyrus, ITG and MOG. Hu et al. (2010) found that both Chinese and English children with DD showed reduced brain activation at the left middle frontal gyrus (MFG), middle temporal gyrus (MTG), angular gyrus and OT area, suggesting language-universal deficits. Feng et al. (2020) found that children with DD in both Chinese and French showed common reduction of brain activation in the left fusiform gyrus and STG.

Even though language-universal deficits in the brain have been suggested in several studies (Feng et al., 2020; Hu et al., 2010; Paulesu et al., 2001), language specificity has been demonstrated as well (Martin et al., 2016; Siok et al., 2004). In a meta-analysis study (Martin et al., 2016), researchers directly compared brain deficits associated with DD between transparent and opaque orthographies and found that functional abnormalities in the brain vary with orthographic depth in alphabetic languages. Specifically, consistent reduction of brain activation was found in a left OT area regardless of orthographic depth including the left ITG, MTG, and inferior occipital gyrus, whereas greater reduction was found in the left fusiform gyrus, left TP and left IFG pars orbitalis in transparent orthographies than in opaque orthographies, and greater reduction in the bilateral intraparietal sulcus, left precuneus and left IFG pars triangularis was found in opaque orthographies than in transparent orthographies. In a recent study on Chinese-English bilingual children with DD, researchers also found both language-universal deficits at the left ITG, the precuneus and dorsal IFG and language-specific deficits at the left IPL for Chinese and the left ventral IFG for English (Cao et al., 2020).These findings suggest that there are both language-universal and language-specific deficits across languages. The language-universal deficits might be related to the causal risk of DD while the language-specific deficits tend to be interpreted as the result of interaction between DD and the specific language system that one studies.

Research on DD in Chinese has revealed different patterns of brain abnormalities from alphabetic languages. Significant alteration in the left MFG or dorsal IFG has been consistently reported in different studies while the alteration in the left TP areas has been reported in only a few studies. Siok et al. (2004, 2008) found that Chinese children with DD showed reduced activation at the left MFG during a homophone judgment task. Cao et al. (2017, 2020) also found reduced activation in the left dorsal IFG in Chinese children with DD compared to age-matched controls and reading-matched controls during an auditory rhyming task and a visual rhyming judgment task. A similar hypoactivation in the left dorsal IFG was also reported in morphological tasks in Chinese (Liu et al., 2012, 2013b). The left dorsal IFG in Cao et al.’s study (2020) was proximal to the left MFG in Siok et al.’s study (2004). In contrast, reduced activation in the left TP area has been found in only a few studies so far (Cao et al., 2017, 2018; Hu et al., 2010).

As a morpho-syllabic language, Chinese characters map to syllables and there are no grapheme-to-phoneme correspondence rules. Each character represents a morpheme and there are many homophones in Chinese; therefore, during Chinese reading, the direct mapping from orthography to semantics plays a very important role. These features of Chinese could help explain why the left TP is less affected and the left dorsal IFG is more affected in Chinese. The left TP has been believed to be essential in rule-based conversion between orthography and phonology and more involved in transparent orthographies than opaque orthographies, while the left dorsal IFG appears to be more involved in whole-word reading in opaque orthographies than in transparent orthographies (Fiez, 2000; Martin et al., 2016; Paulesu et al., 2000). It has also been found that the left TP is more involved in English reading and the left dorsal IFG is more involved in Chinese reading (Bolger et al., 2005; Tan et al., 2005). Therefore, the greater deficit in the left dorsal IFG than in the left TP areas in Chinese is consistent with the features of Chinese. Chinese characters also have complex visuo-spatial configurations, which require an increasing involvement of visual-orthographic analysis at the bilateral precuneus (Cao et al., 2013). That explains the fact that the precuneus showed consistent deficit in Chinese DD (Cao et al., 2018, 2020). Ziegler (2006) argued that Chinese readers with DD should share a common neurocognitive deficit that DD readers have in alphabetic languages, because Chinese reading is essentially the process of mapping from orthography to phonology and meaning as alphabetic languages do and Chinese readers with DD also show phonological deficits (Goswami et al., 2011; Hu et al., 2010). However, despite a universal phonological deficit across languages, different parts of the brain might be affected, because different strategies might be used when mapping orthography to phonology depending on the features of the language.

A large number of studies have been conducted in investigating structural alterations associated with DD; however, none of them has taken language difference into account. In a meta-analysis, Linkersdorfer et al. (2012) gathered nine voxel-based morphometry (VBM) studies in alphabetic languages and found consistent grey matter reduction in the left OT area and TP area, but failed to find reliable evidence in the left IFG. In another two meta-analysis studies in alphabetic languages, researchers only found consistent grey matter reduction in the left TP area but failed to find reduction in the OT area (McGrath & Stoodley, 2019; Richlan et al., 2013). These three meta-analytic studies echo findings from functional studies by showing abnormal brain structures within the classic reading network. However, studies also showed abnormal brain structures outside the reading network. For example, studies have found reduced grey matter volume (GMV) at the left putamen in Chinese children with DD (Wang et al., 2019), at the cerebellum in Spanish (Adrian-Ventura et al., 2020), Italian (Brambati et al., 2004) and English (Brown et al., 2001), at the thalamus in English (Brown et al., 2001), French, German and Polish (Jednorog et al., 2015), and at the caudate nucleus in English (Brown et al., 2001; Jagger-Rickels et al., 2018). In summary, previous inconsistent findings in brain structure might be due to the lack of differentiation in participants’ language. It is important to differentiate language-universal structural alterations as a core deficit which might be related to the cause of DD and language-specific structural alterations as a consequence of being DD in a specific language. While brain structural deficits may cause reading disability, learning experience may also shape brain development. Learning a specific language with DD may affect brain development in language-specific regions (Mechelli et al., 2004).

DD is associated with altered brain structure and function, but very few studies have investigated whether brain structural alterations and brain functional alterations are consistent or inconsistent. In a study by Siok et al. (2008), researchers examined both structural and functional alterations in Chinese children with DD, and found reduced GMV and brain activation in the left MFG, which underscores the association between the left MFG and DD in Chinese. Another study located a key region in the left IPL, which showed reduced GMV and activation in English-speaking readers with DD (Hoeft et al., 2007). There is a paucity of research to examine brain regions that show increased GMV but decreased brain function or vice versa, and neither is there discussion about the neurocognitive implications of such patterns. Simultaneously considering structural and functional abnormalities with a focus on cross-linguistic comparison would provide a comprehensive perspective to understand the neural mechanisms of DD.

In this meta-analysis study, we aimed to explore how structural and functional impairment of DD converge or diverge and whether this pattern is similar or different across writing systems. We expected to find brain regions that show decreased brain structure and function, indicating insufficient neuronal resources for certain cognitive computations. For regions that show increased brain structure and function, we believe they develop to an unusually high degree for compensation. For brain regions with increased structure but decreased function or decreased structure and increased function, it may be due to brain structures receiving inhibitory input from other regions. We also expected to find language-universal as well as language-specific neurological abnormalities. For language-universal deficits, we tend to believe that they are related to the cause of DD, while the language-specific deficits tend to be consequences of DD in different languages.

## Methods

### Literature retrieval and data extraction

We searched in “PubMed” (http://www.pubmed.org) and “Web of science” using a combination of “dyslexia”, “reading disorder”, “reading impairment” or “reading disability” and “fMRI”, “PET”, “voxel-based morphometry”, “VBM” or “neuroimaging” as key words for neuroimaging studies published from January 1986 to January 2020. Additionally, we manually added studies by checking the references of the selected papers that were missed in the search. The inclusion criteria were: (1) PET, fMRI, voxel-based morphometry (VBM) studies or structural studies using a volumetric FreeSurfer pipeline, (2) whole-brain results were reported, (3) direct group comparisons between readers with DD and age control readers were reported, (4) coordinates were reported in Talairach or MNI stereotactic space, and (5) studies on DD in the first language. The exclusion criteria were (1) studies with only ROI analysis, (2) resting-state studies, (3) studies that only included readers with DD or did not report group differences, (4) studies with direct group comparisons only between readers with DD and reading level control readers, (5) studies on children at risk for dyslexia, and (6) studies focused on non-linguistic tasks (Evans et al., 2014a; Margolis et al., 2019; Menghini et al., 2006; Yang et al., 2013). Finally, 119 experiments from 110 papers were included in this meta-analysis comprising 92 brain functional experiments (from 87 papers) and 27 brain structural experiments (from 23 papers) (see Table 1, Table 2 and Figure 1 for detail). From the original publications, we extracted peak coordinates, where there is a significant difference between controls and individuals with DD either in brain activation or regional GMV. We also extracted effect sizes and other information from the publications.

**Figure 1.**
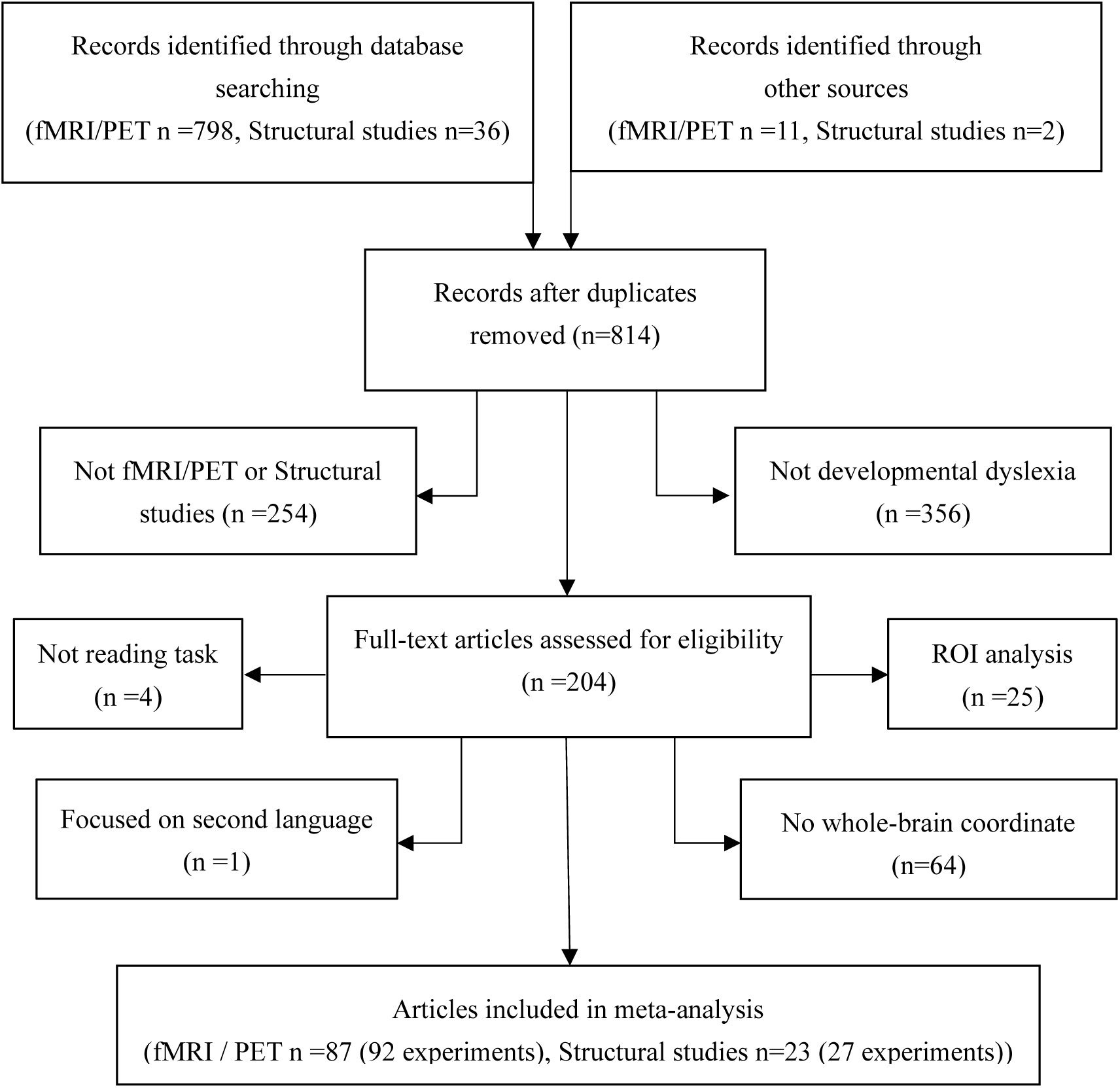
PRISMA flowchart of the selection process for included articles.

**Table 1.**
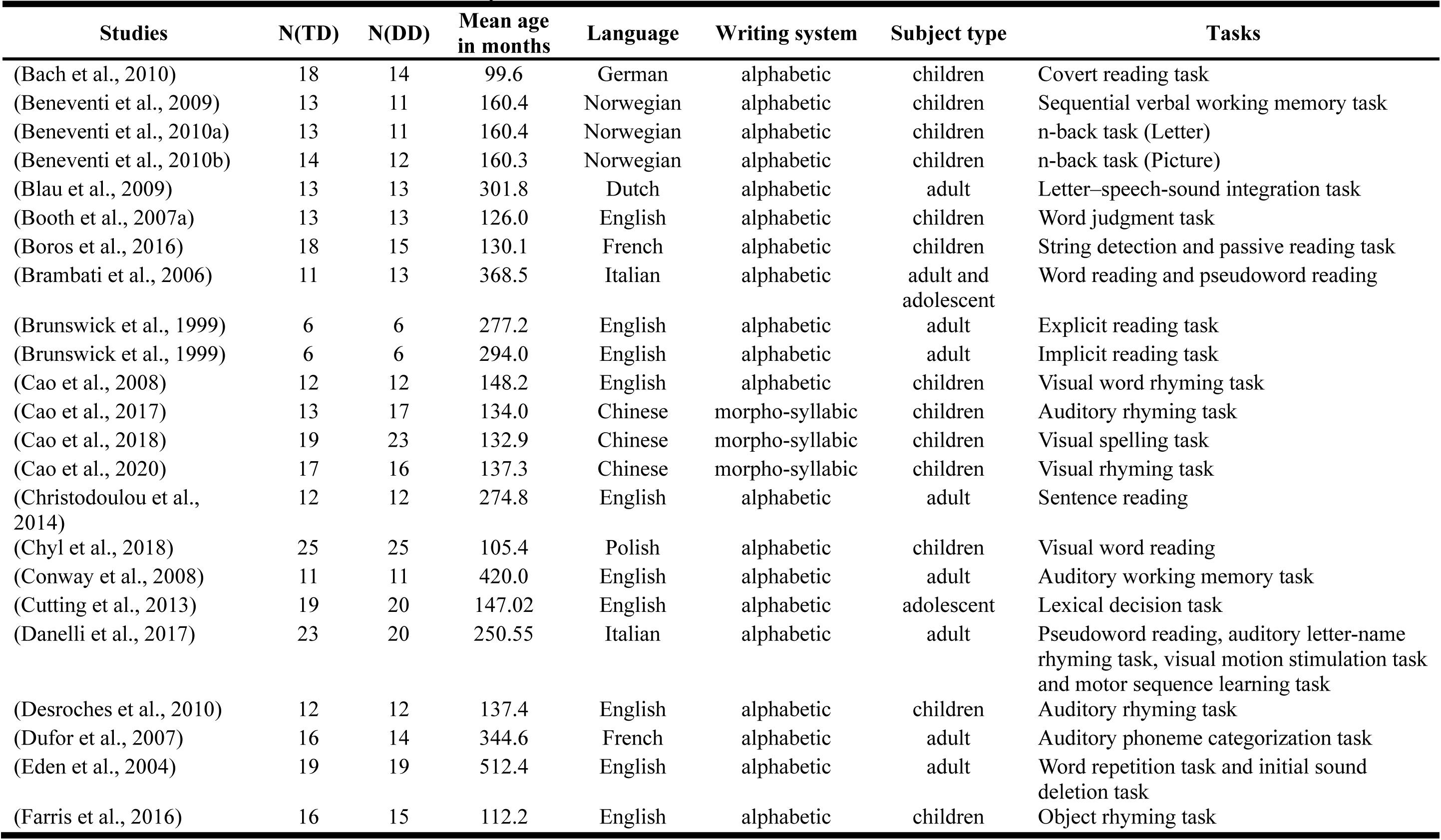
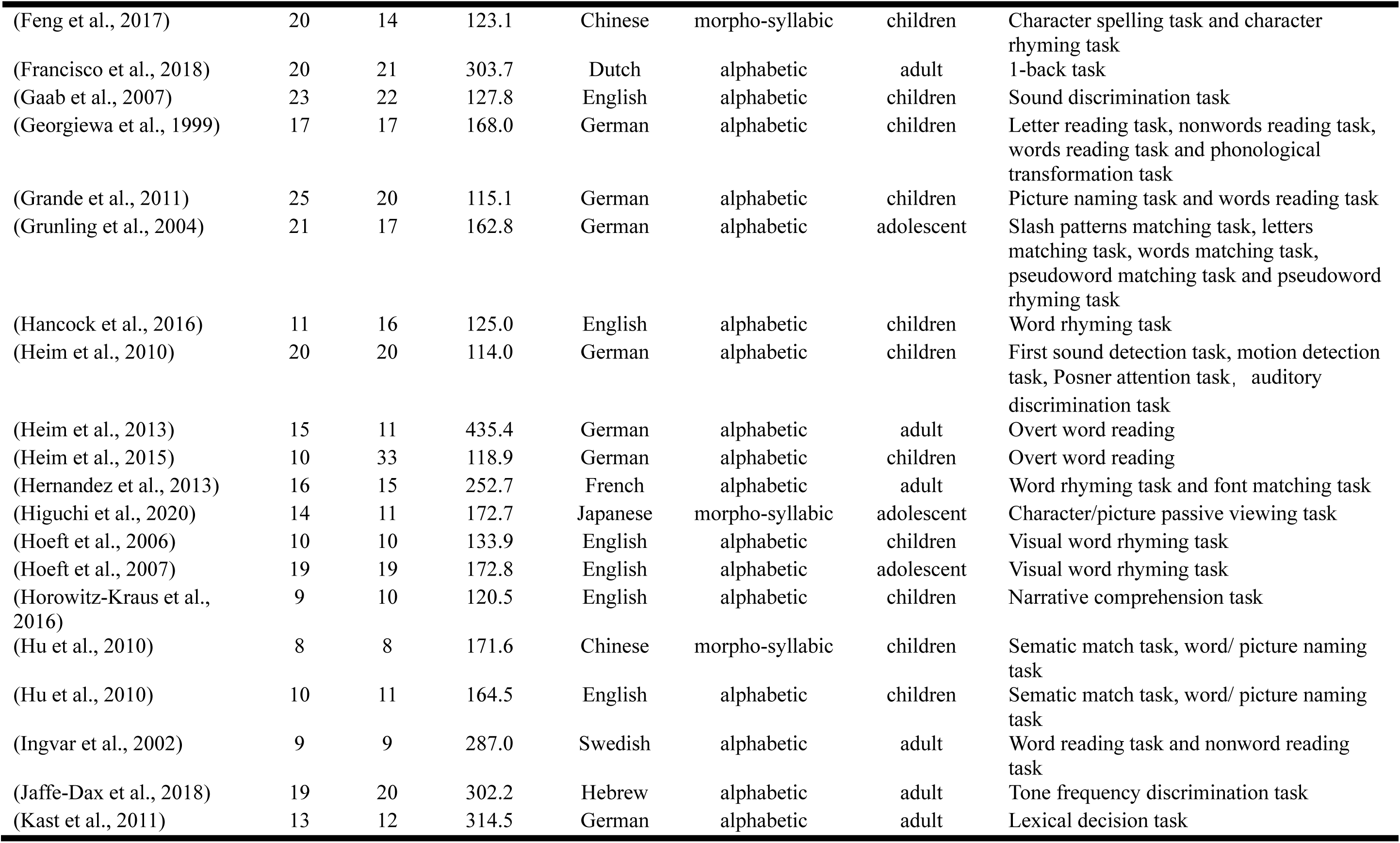
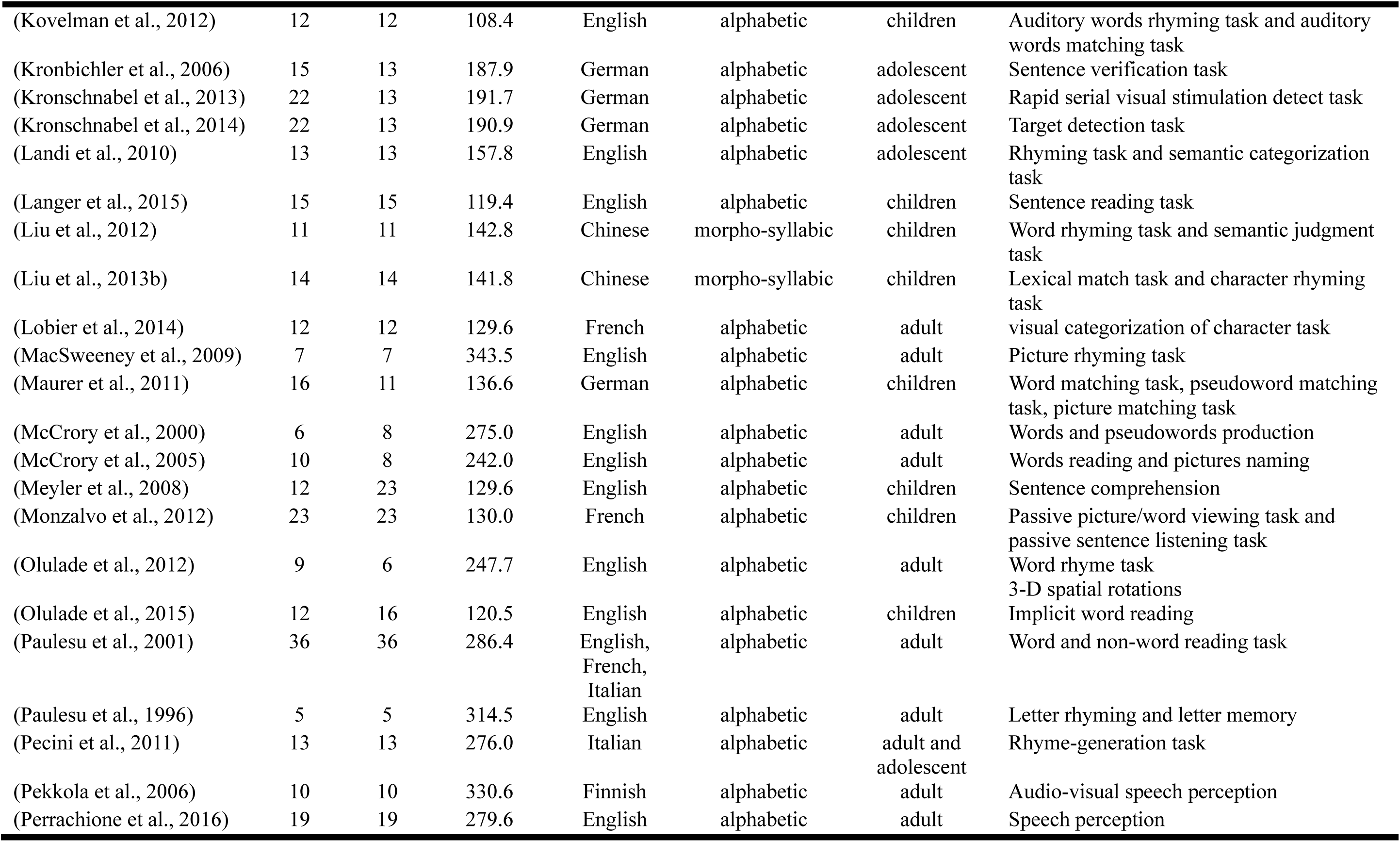
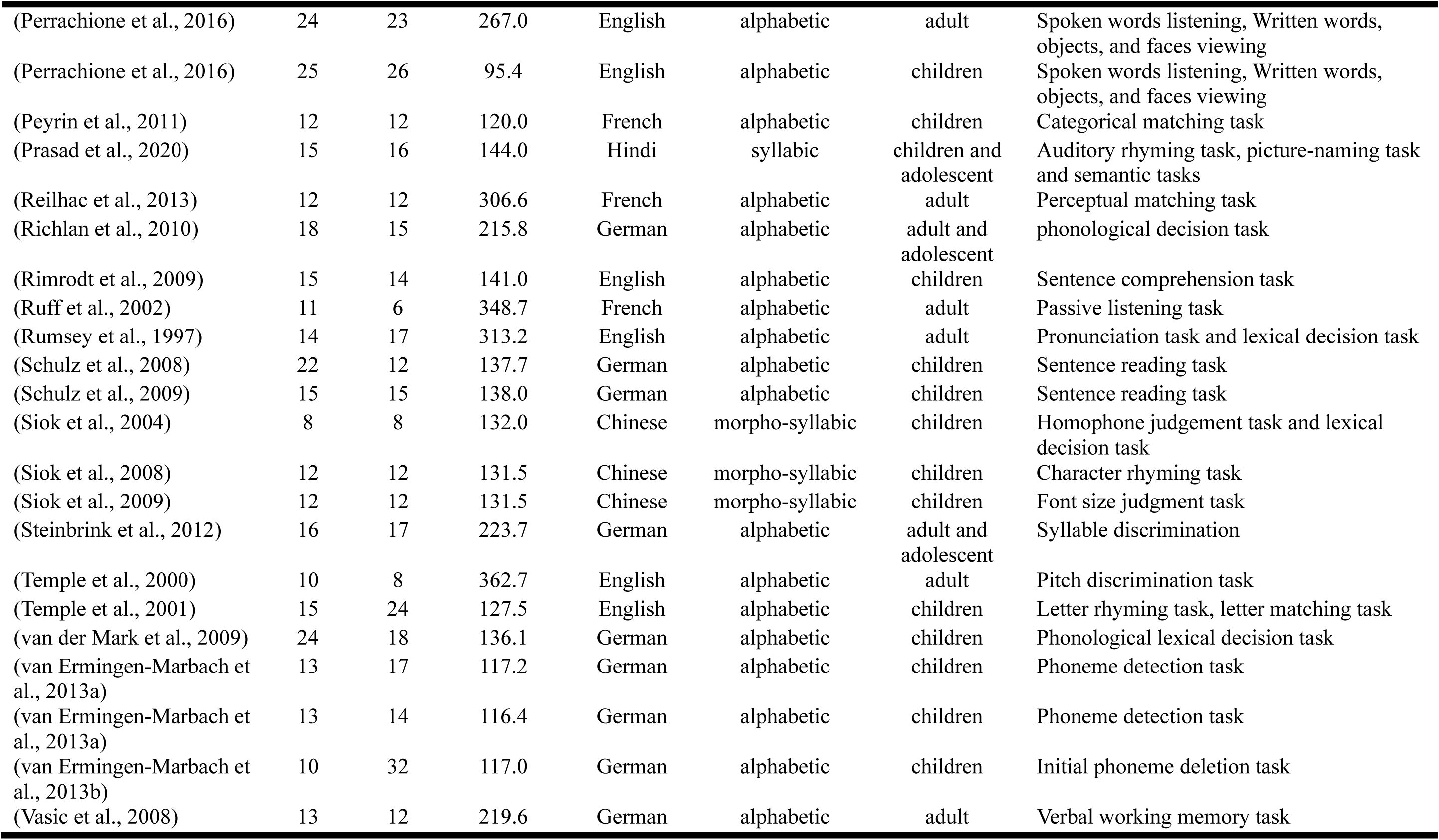
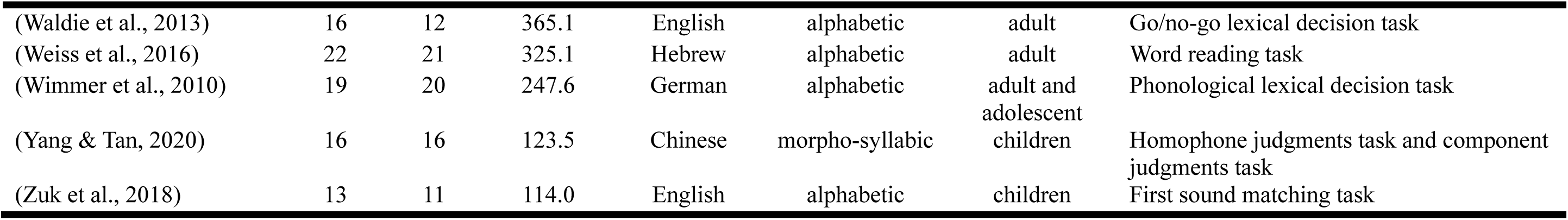
functional studies included in the meta-analysis.

**Table 2.**
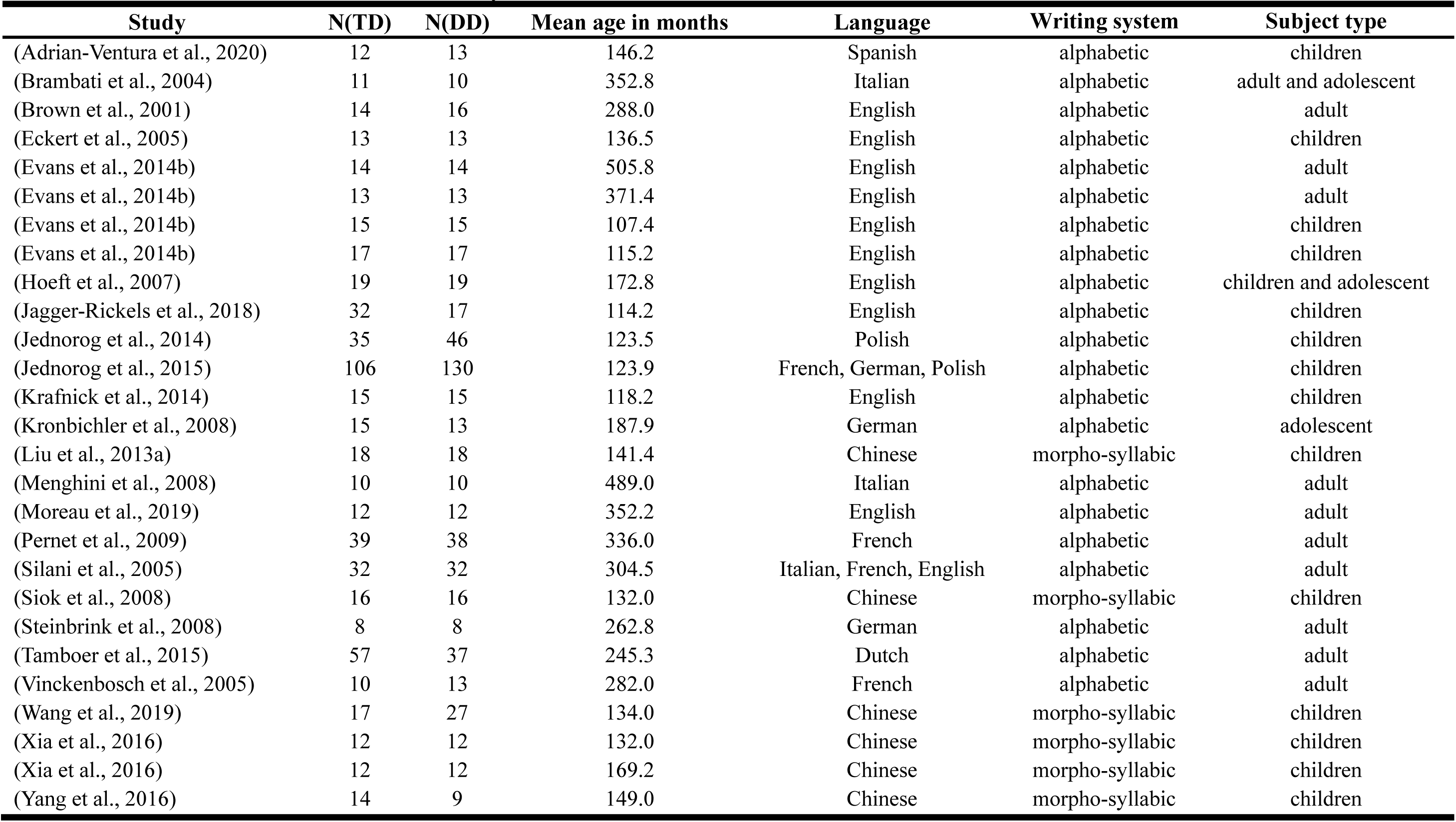
Structural studies included in the meta-analysis.

### Voxel-wise meta-analysis

After data acquisition, we conducted a voxel-wise meta-analysis using the anisotropic effect-size version of Signed Differential Mapping software (AES-SDM version 5.14, see http://www.sdmproject.com) separately for functional studies and structural studies. Unlike other coordinate-based meta-analysis methods such as Activation likelihood estimation (ALE) or Multilevel peak Kernel density analysis (MKDA), AES-SDM combined the peak coordinates with the statistical parameter maps to increase the sensitivity of the analysis (Radua et al., 2012a). Data were first preprocessed with the statistical parameter maps and the peak coordinates were convolved with a fully anisotropy un-normalized Gaussian kernel (ɑ=1) (full width at half maximum = 20 mm) to recreate the effect size map and the corresponding variance map for each study (Radua et al., 2012a, 2014). Then, a random-effect model was set up to calculate the differences between the DD group and the control group. Five hundred permutations were performed to ensure the stability of the analysis. Finally, the results of the standard meta-analysis were thresholded at peak height of the mean effect size SDM-Z=1, uncorrected p=.005 at the voxel level and 150 voxels at the cluster level, which is stricter than the advised threshold by Radua et al. (Radua et al., 2012a) (peak height SDM-Z=1, uncorrected p=.005 at the voxel level and 10 voxels at the cluster level) in order to avoid false-positive results.

To test the stability of the meta-analysis results, we conducted a whole-brain jack-knife sensitivity analysis. The standard meta-analysis was repeated n times (n=92 for functional studies, n=27 for structural studies) but leaving out one study each time, to determine whether the results remained significant.

### Multimodal meta-analysis

Because we were interested in the convergence between functional deficits and structural deficits, a multimodal meta-analysis was conducted, which provided an efficient way to combine two meta-analyses in different modalities. The union probabilities of the meta-analytical maps of functional studies and structural studies were estimated and then thresholded at the peak height p=0.00025, with a voxel level uncorrected p=0.0025 and 150 voxels at the cluster level, which was stricter than the one suggested by Radua et al. (2012b, 2013) (peak height p=0.00025, with a voxel level uncorrected p=0.0025 and 10 voxels at cluster level).

### Subgroup meta-analysis

In order to explore the language effect, we conducted a subgroup meta-analysis. According to the native language of the participants, the functional and structural studies were subdivided into an alphabetic language group in which writing symbols represent phonemes, and a morpho-syllabic language group in which each writing symbol represents a morpheme with a syllable, which resulted in 79 functional and 21 structural studies for alphabetic languages and 12 functional (including a Japanese study focused on Kanji) and 6 structural studies for morpho-syllabic languages. First, a standard functional and structural meta-analysis was conducted for alphabetic languages and morpho-syllabic languages, separately. Then multimodal meta-analysis was conducted to find the convergence between functional and structural deficits separately for each language group. The threshold for the separate structural and functional maps and multimodal analysis was the same as mentioned above. Next, we conducted a direct comparison between the alphabetic language and morpho-syllabic language group for functional studies and structural studies separately to find the difference between the two language groups. The threshold was set at peak height SDM-Z=1, voxel level uncorrected p=.005 and 150 voxels at the cluster level.

### Confirmation study

Because there were many more functional studies included in the alphabetic group than in the morpho-syllabic group, and most of the morpho-syllabic studies focused on children, the language difference may be due to these differences in the two groups of studies. To avoid influences of these factors, we selected ten English studies (Booth et al., 2007a; Cao et al., 2008; Farris et al., 2016; Hancock et al., 2016; Hu et al., 2010; Langer et al., 2015; Meyler et al., 2008; Olulade et al., 2015; Rimrodt et al., 2009; Temple et al., 2001) and ten Chinese studies (Cao et al., 2018, 2020; Feng et al., 2017; Hu et al., 2010; Liu et al., 2012, 2013b; Siok et al., 2004, 2008, 2009; Yang & Tan, 2020) for further confirmation analysis. The two subgroups were matched on participants’ age (mean age = 10.95 years for English studies, mean age = 11.40 years for Chinese studies) and task (visual word tasks). Then, we conducted a direct comparison between the Chinese studies and English studies. The threshold was set at peak height SDM-Z=1 and voxel level uncorrected p=.005 with 150 voxels at the cluster level.

## Results

### Description of the included studies

For the functional studies, a total of 2746 participants (controls:1377, DD:1369) were included, and the mean age was 16.55 years for controls and 16.23 years for participants with DD. For the structural studies, a total of 1183 participants (controls:588, DD:595) were included, and the mean age was 16.88 years for controls and 16.97 years for participants with DD.

### Meta-analysis results

#### Functional impairment associated with DD across all languages

In the meta-analysis of all functional studies, hypoactivation in DD was found in a large cluster peaked at the left IPL which extended to the inferior frontal cortex, occipitotemporal cortex and cerebellum, and a cluster peaked at the right MOG (Table S1 and Figure S1). Hyperactivation in DD was found in the right cerebellum, right precentral/postcentral gyrus and bilateral caudate nucleus. The jack-knife sensitivity analysis showed that all results reported above were replicable (Table S1).

#### Structural impairment associated with DD across all languages

In the meta-analysis of all structural studies, readers with DD showed a decrease in GMV in the left inferior frontal cortex and right STG (Table S2 and Figure S2). In contrast, readers with DD showed an increase in GMV in the right MTG and left IPL. The jack-knife sensitivity analysis showed that all results reported were replicable (Table S2).

#### Multimodal analysis across all languages

As shown in Table 3 and Figure 2, decreased GMV and hypoactivation in DD were found in the left IFG, and left TP which peaked at the left STG and extended to the left IFG and precentral gyrus. Increased GMV and hyperactivation in DD were found in the right anterior MTG/ITG. Increased GMV and hypoactivation in DD were found in the left IPL and left cerebellum which extended to FG. Decreased GMV and hyperactivation in DD were found in bilateral caudate.

**Figure 2.**
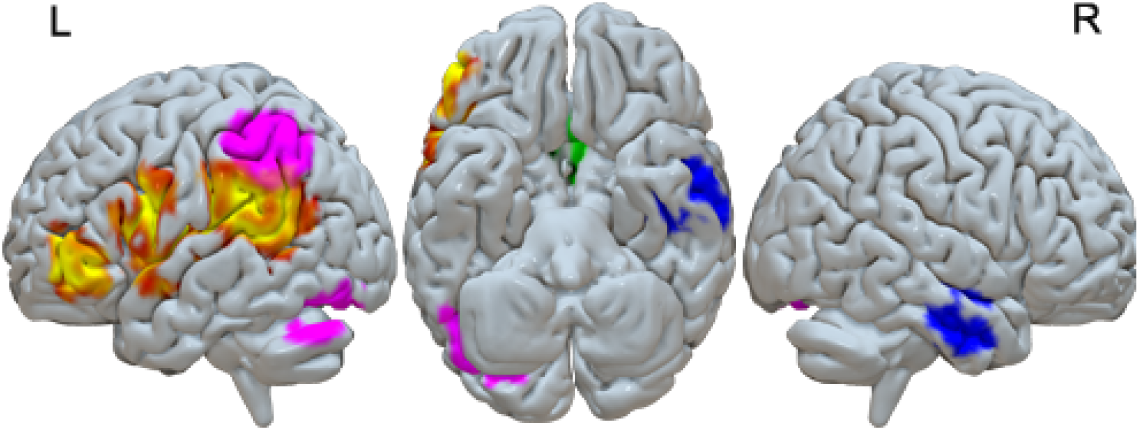
Structural and functional deficits in DD across all languages (red-yellow: decreases in DD in both structural and functional, blue: increases in DD in both structural and functional studies, purple: increases in DD in structural studies and decreases in DD in functional studies, green: decreases in DD in structural studies and increases in DD in functional studies).

**Table 3.** Multi-modal structural and functional abnormalities in individuals with DD across all languages.

| Regions | MNI coordinate | Voxels | Cluster breakdown (Voxels) |
| --- | --- | --- | --- |
| *Decreased GMV and hypoactivation in DD* |  |  |  |
| Left STG | -54,-34,20 | 3597 | Left rolandic operculum, BA 48 (388)<br>Left STG, BA 48 (313)<br>Left supramarginal gyrus, BA 48 (282)<br>Left STG, BA 42 (262)<br>Left insula, BA 48 (240)<br>Left MTG, BA 21 (170)<br>Left IFG, opercular part, BA 48 (152) |
| Left IFG | -46,42,0 | 354 |  |
| *Increased GMV and hypoactivation in DD* |  |  |  |
| Left IPL | -40,-38,38 | 1439 | Left IPL, BA 40 (822) |
| Left cerebellum | -42,-68,-24 | 709 | Left fusiform gyrus, BA 19 (196)<br>Left cerebellum, crus I, BA 19 (162) |
| *Decreased GMV and hyperactivation in DD* |  |  |  |
| Right caudate | 8,8,10 | 559 | Right anterior thalamic projections (204)<br>Right caudate nucleus (180) |
| Left caudate | -18,10,10 | 519 | Left anterior thalamic projections (323) |
| *Increased GMV and hyperactivation in DD* |  |  |  |
| Right MTG | 48,-8,-24 | 669 | Right inferior network (238)<br>Right ITG, BA 20 (166)<br>Right MTG, BA 21 (151) |

We also found brain regions that had normal structure but altered function. We found normal brain structure but reduced brain activation at the right MOG and a large cluster peaked at left supramarginal gyrus, which extended to the left ITG, MTG and IFG. We found normal brain structure but increased activation at the right cerebellum and right precentral gyrus (Figure S7 and Table S7). There are also regions that had altered structure but normal function. We found normal brain activation but reduced GMV at the left IFG and right STG, as well as normal brain function but increased GMV at the right MTG (Figure S8 and Table S8).

#### Subgroup analysis results – comparison between alphabetic and morpho-syllabic languages

For the direct comparison between the morpho-syllabic and alphabetic languages in functional studies, we found greater reduction of brain activation in alphabetic languages than in morpho-syllabic languages in the right STG, left MTG and left fusiform gyrus. We found greater reduction of brain activation in morpho-syllabic languages than in alphabetic languages in the left IFG and greater increase of brain activation in DD in morpho-syllabic languages than in alphabetic languages in the right precentral gyrus (Table 4, Figure 3).

**Figure 3.**
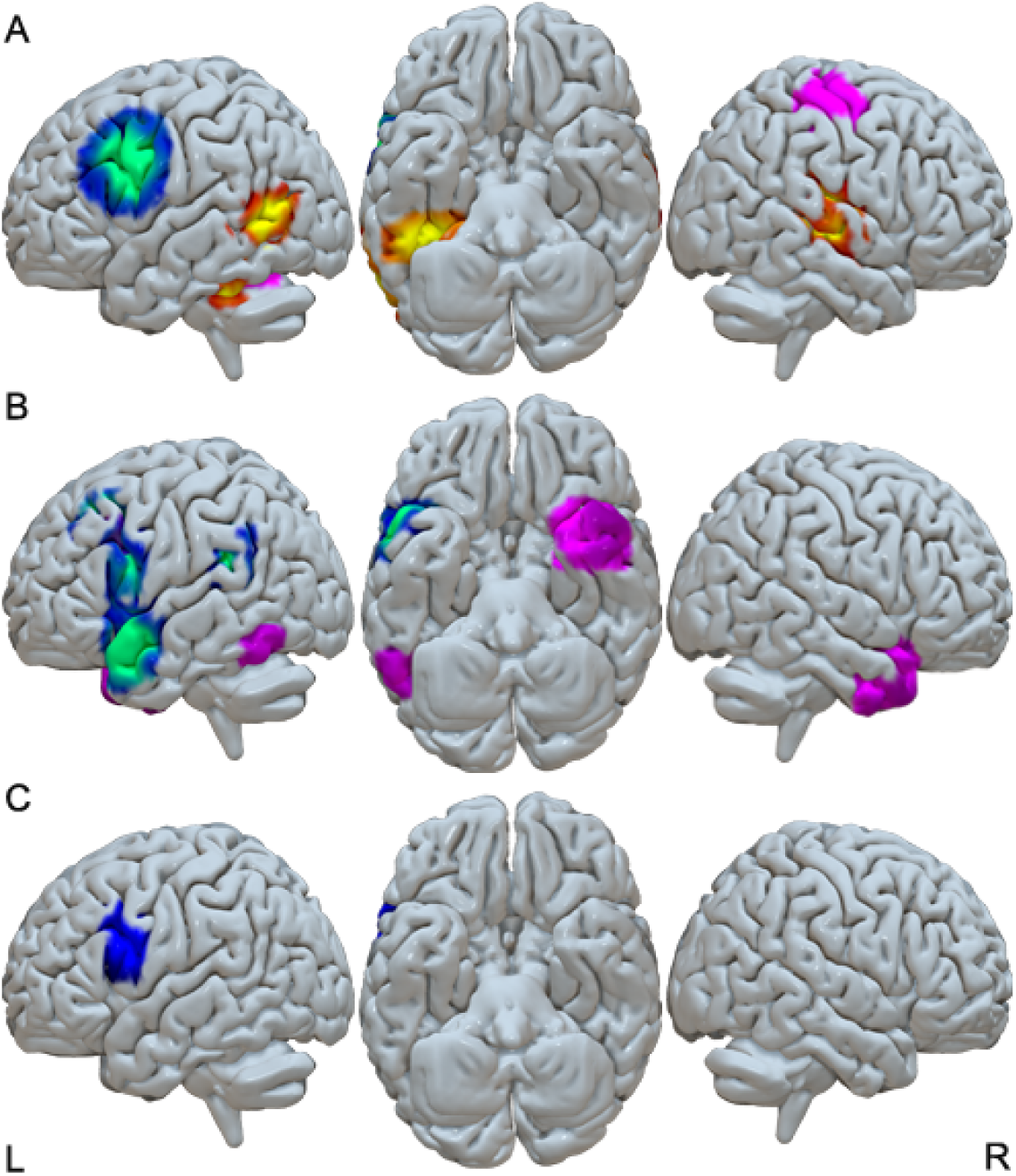
Direct comparison between alphabetic languages and morpho-syllabic languages in structural and functional deficits. A, Language differences in functional deficits. B, Language differences in structural deficits. C, Conjunction of language differences between functional and structural deficits, which is due to greater reduction in brain activation and GMV in morpho-syllabic languages than in alphabetic languages in the left dorsal IFG. (red-yellow: greater decreases in DD in alphabetic languages than in morpho-syllabic languages; blue-green: greater decreases in DD in morpho-syllabic languages than in alphabetic languages; purple: greater increases in DD in morpho-syllabic languages than in alphabetic languages).

**Table 4.** Direct comparison between alphabetic languages and morpho-syllabic languages in functional studies.

| Regions | MNI coordinate | SDM-Z | P | Voxels | Cluster breakdown (Voxels) |
| --- | --- | --- | --- | --- | --- |
| *Hypoactivation in DD* |  |  |  |  |  |
| *Alphabetic languages > Morpho-syllabic languages* |  |  |  |  |  |
| Right STG | 60,-18,4 | 1.059 | 0.0000 | 2448 | Right STG, BA 22 (436)<br>Corpus callosum (381)<br>Right STG, BA 48 (287)<br>Right insula, BA 48 (271)<br>Right rolandic operculum, BA 48 (261) |
| Left fusiform gyrus | -38,-42,-22 | 1.215 | 0.0000 | 846 | Left fusiform gyrus, BA 37 (187)<br>Left ITG, BA 20 (183) |
| Left MTG | -50,-66,8 | 1.121 | 0.0000 | 732 | Left MTG, BA 37 (396) |
| *Morpho-syllabic languages > Alphabetic languages* |  |  |  |  |  |
| Left IFG | -48,8,28 | -4.282 | 0.0000 | 2179 | Left precentral gyrus, BA 6 (524)<br>Left IFG, opercular part, BA 44 (284)<br>Left precentral gyrus, BA 44 (191)<br>Left IFG, triangular part, BA 48 (174)<br>Corpus callosum (167) |
| *Hyperactivation in DD* |  |  |  |  |  |
| *Morpho-syllabic languages > Alphabetic languages* |  |  |  |  |  |
| Right precentral gyrus | 40,-20,54 | -2.262 | 0.0000 | 1518 | Right precentral gyrus, BA 6 (525)<br>Right precentral gyrus, BA 4 (306)<br>Right postcentral gyrus, BA 3 (286)<br>Right postcentral gyrus, BA 4 (171) |

For the direct comparison between the morpho-syllabic and alphabetic languages in structural studies, we found greater reduction of GMV in DD in morpho-syllabic languages than in alphabetic languages in the left STG, left IFG, left MFG, left supramarginal gyrus, left superior occipital gyrus (SOG) and left insula. We also found greater increase of GMV in DD in morpho-syllabic languages than in alphabetic languages in the right STG and left ITG (Table 5, Figure 3). We found no regions that showed greater GMV changes in alphabetic languages than in morpho-syllabic languages.

**Table 5.** Direct comparison between alphabetic languages and morpho-syllabic languages in structural studies.

| Regions | MNI coordinate | SDM-Z | P | Voxels | Cluster breakdown (Voxels) |
| --- | --- | --- | --- | --- | --- |
| *Decreased GMV in DD* |  |  |  |  |  |
| *Morpho-syllabic languages > Alphabetic languages* |  |  |  |  |  |
| Left STG | -50,10,-18 | -3.139 | 0.0000 | 1218 | Left STG, BA 38 (322) |
| Left IFG | -52,10,24 | -2.253 | 0.0008 | 409 |  |
| Left MFG | -28,16,44 | -2.888 | 0.0001 | 346 |  |
| Left supramarginal gyrus | -48,-44,24 | -2.761 | 0.0001 | 344 |  |
| Left SOG | -20,-72,20 | -2.911 | 0.0001 | 287 | Corpus callosum (187) |
| Left insula | -32,14,8 | -2.046 | 0.0015 | 179 |  |
| *Increased GMV in DD* |  |  |  |  |  |
| *Morpho-syllabic languages > Alphabetic languages* |  |  |  |  |  |
| Right STG | 34,6,-28 | -1.893 | 0.0001 | 1397 | Right STG, BA 38 (246)<br>Right inferior network (239) |
| Left ITG | -54,-58,-16 | -1.120 | 0.0018 | 161 |  |

To identify the common language differences across both structural and functional studies, we conducted a conjunction analysis of the thresholded language difference maps. This produced an overlap of 387 voxels in the left IFG, which peaked at (-48, 8, 26), indicating greater reduction of both GMV and brain activation in morpho-syllabic languages than in alphabetic languages (Figure 3).

#### Subgroup analysis results – multimodal analysis

Multimodal meta-analysis in alphabetic languages showed that decreased GMV and hypoactivation in DD were found in the bilateral TP and left IFG; no regions showed increased GMV and hyperactivation; increased GMV and hypoactivation in DD were found in left IPL and left cerebellum; decreased GMV and hyperactivation in DD were found in bilateral caudate and right cerebellum (Table 6, Figure 4). Multimodal meta-analysis in morpho-syllabic languages showed that decreased GMV and hypoactivation in DD were found in the left TP and left IFG; increased GMV and hyperactivation in DD were found in the right MTG; decreased GMV and hyperactivation in DD were found in left STG; no regions showed increased GMV and hypoactivation (Table 7, Figure 4).

**Figure 4.**
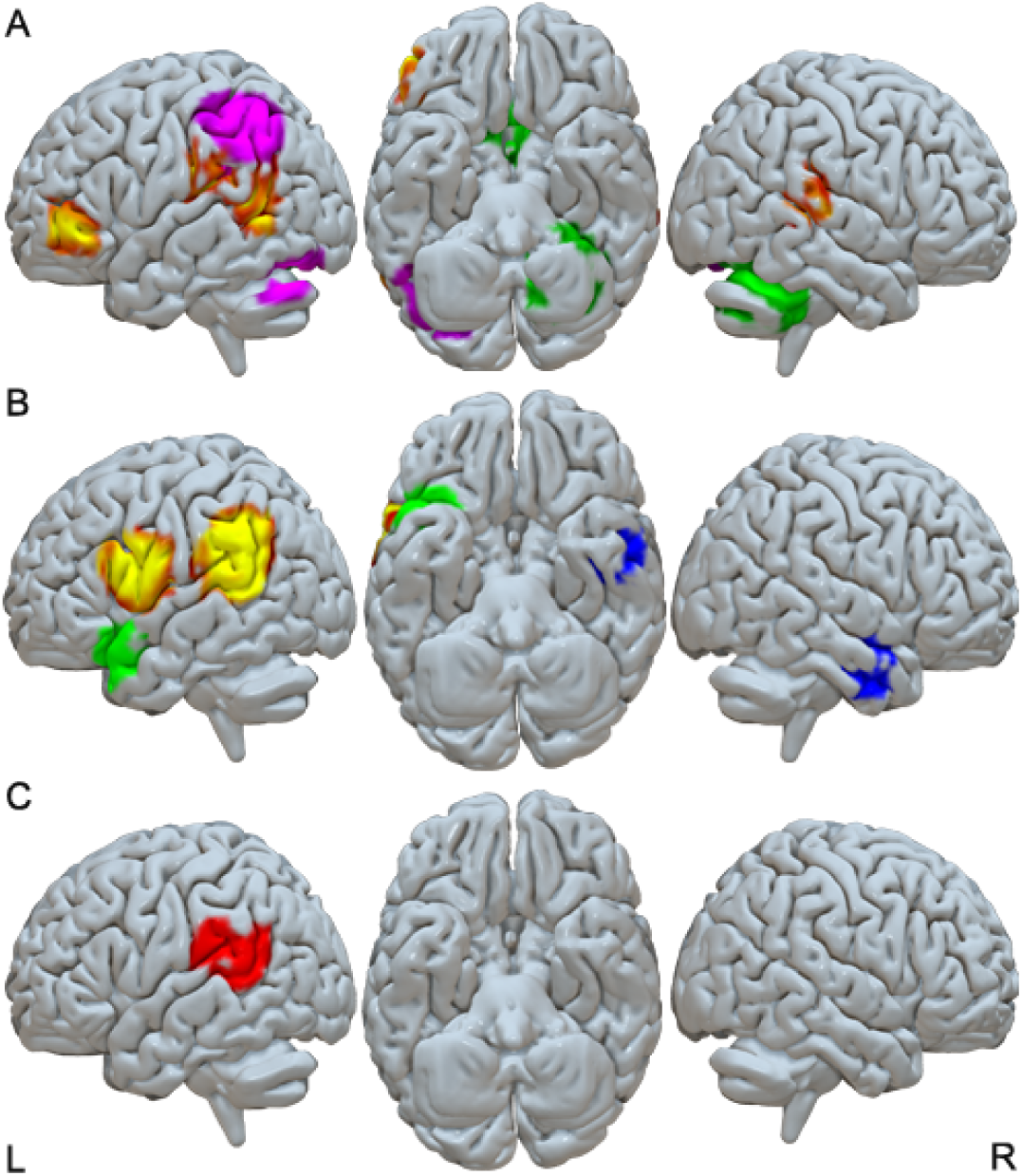
Structural and functional deficits in DD for alphabetic languages and morpho-syllabic languages. A, Structural and functional deficits in DD for alphabetic languages. B, Structural and functional deficits in DD for morpho-syllabic languages. C, Conjunction of structural and functional deficits in alphabetic and morpho-syllabic languages, which is driven by decreased GMV and brain activation in the left STG in both alphabetic and morpho-syllabic languages. (red-yellow: decreases in DD in both structural and functional studies, blue: increases in DD in both structural and functional studies, purple: increases in DD in structural studies and decreases in functional studies, green: decreases in DD in structural studies and increases in functional studies).

**Table 6.** Multi-modal structural and functional abnormalities in individuals with DD in alphabetic languages.

| Regions | MNI coordinate | Voxels | Cluster breakdown (Voxels) |
| --- | --- | --- | --- |
| *Decreases of GMV and hypoactivation in DD* |  |  |  |
| Left supramarginal gyrus | -50,-30,20 | 1087 | Left MTG, BA 21 (236) |
| Right STG | 62,-32,14 | 328 |  |
| Left IFG | -46,42,0 | 240 |  |
| *Increases of GMV and hypoactivation in DD* |  |  |  |
| Left IPL | -46,-40,38 | 1692 | Left IPL, BA 40 (942) |
| Left cerebellum | -40,-70,-24 | 433 | Left cerebellum, crus I, BA 19 (154) |
| *Decreases of GMV and hyperactivation in DD* |  |  |  |
| Right cerebellum | 28,-52,-34 | 1313 | Right cerebellum, lobule VI, BA 37 (75)<br>Middle cerebellar peduncles (272)<br>Right cerebellum, lobule VI, BA 19 (207)<br>Right cerebellum, crus I (155) |
| Right caudate | 8,8,12 | 600 | Right anterior thalamic projections (214)<br>Right caudate nucleus (189) |
| Left caudate | -16,8,14 | 595 | Left anterior thalamic projections (353) |

**Table 7.**
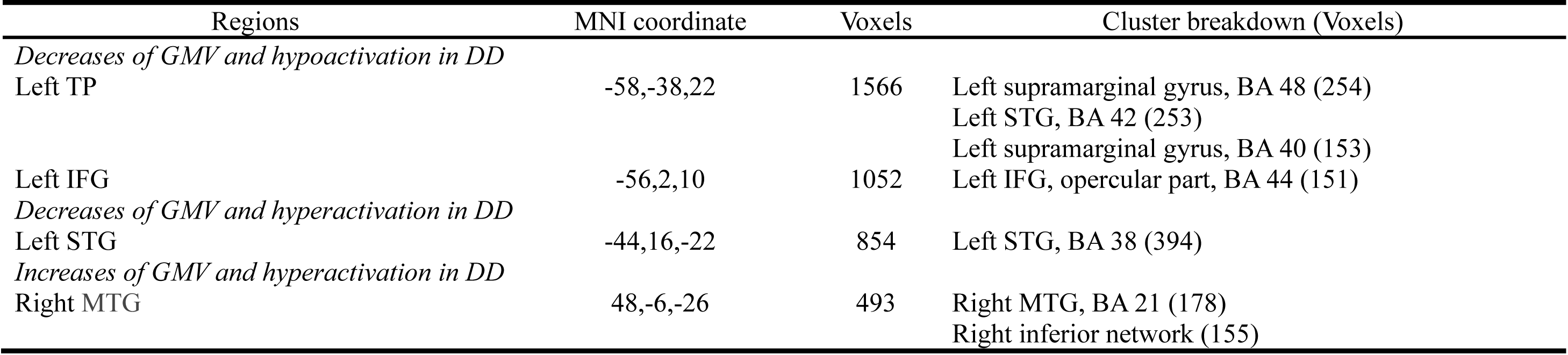
Multimodal structural and functional abnormalities in individuals with DD in morpho-syllabic languages.

To identify the common structural and functional deficit in alphabetic languages and morpho-syllabic languages, we conducted a conjunction analysis of the thresholded multimodal maps of the two types of writing systems. This procedure produced an overlap of 482 voxels in the left STG, which peaked at (-56, -32, 20), indicating shared reduction of GMV and hypoactivation in both types of writing systems (Figure 4).

#### Subgroup analysis results – confirmation analysis

Confirmation analysis of two well-matched subgroups showed greater reduction of brain activation in DD in English than in Chinese in the left ITG and left MOG and greater reduction of brain activation in DD in Chinese than in English in the left precentral gyrus. Greater increase of brain activation in English than in Chinese was found in the right supramarginal gyrus, right hippocampus, and right supplementary motor area; greater increase of brain activation in Chinese than in English was found in the right precentral gyrus (Table S9, Figure S9). Conjunction analysis of the difference maps between alphabetic and morpho-syllabic languages and the difference maps between English and Chinese showed consistent greater reduction of brain activation in DD in alphabetic languages/English than in morpho-syllabic languages/Chinese in the left ITG (-50, -42, -24) and left MTG (-50, -66, 8) with a cluster size of 76 voxels and 147 voxels, respectively; there was consistent greater reduction of brain activation in DD in morpho-syllabic languages/Chinese than in alphabetic languages/English in the left IFG (-46, 8, 30) with a cluster size of 1999 voxels. Consistent greater increase of brain activation in morpho-syllabic languages/Chinese than in alphabetic languages/English was also found in the right precentral gyrus (42, -20, 52), and the cluster size was 791 voxels. The results from the well-matched subgroup analysis confirmed findings from the comparison between alphabetic languages and morpho-syllabic languages.

## Discussion

In this meta-analysis study, we examined the relationship between structural and functional deficits associated with DD as well as whether the deficits are consistent across languages. We found that readers with DD showed both GMV reduction and functional hypoactivation in the left TP and inferior frontal cortex, among which, the left STG was a shared impairment across all languages, and the left IFG showed a greater impairment in morpho-syllabic languages than in alphabetic languages, suggesting both language-universal and language-specific deficits in the brain. We also found GMV increase and functional hyperactivation in the right anterior MTG/ITG region across all languages; however, conjunction analysis between morpho-syllabic languages and alphabetic languages did not reveal any overlap. In addition to the consistent structural and functional alterations, we also detected inconsistent structural and functional alterations. Individuals with DD showed increased GMV and hypoactivation in the left IPL and left cerebellum, and decreased GMV and hyperactivation in the bilateral caudate. However, when we subdivided the data into alphabetic and morpho-syllabic languages, these inconsistent structural and functional changes only existed in alphabetic languages.

### Convergent structural and functional impairment across writing systems

Across writing systems, convergent structural and functional deficits were found around the left perisylvian region (i.e., left IFG, left supramarginal gyrus and left STG) due to reduced GMV and brain activation. The left perisylvian region is a very important component in the language network, which shows structural and functional abnormality in individuals with DD as suggested in previous meta-analysis studies (Linkersdorfer et al., 2012; Maisog et al., 2008; McGrath & Stoodley, 2019; Paulesu et al., 2014; Richlan et al., 2013). The left STG and the dorsal IFG are two core regions in the language model proposed by Friederici (2012), and by Hickok and Poeppel (2007). These two brain regions are involved in phonological processing during both spoken language processing and reading (Bolger et al., 2005; Enge et al., 2020; Tan et al., 2005). Furthermore, proficient reading is characterized by convergence between speech and print at these two regions regardless of languages, because multivariate brain activity patterns are similar for speech and print at these regions (Chyl et al., 2021). These findings suggest that the reading network may develop based on the built-in language circuit, as reading is a skill that humans acquire too late in the course of human evolution to have a brain network dedicated to it. Our finding suggests that dyslexia is associated with structural and functional abnormalities of the language network regardless of language. Recently, a growing number of studies have investigated early signs of dyslexia before the onset of reading and found that structural and functional deficits in the left TP area and left inferior frontal cortex appear before reading onset (Clark et al., 2014; Hosseini et al., 2013; Plewko et al., 2018; Raschle et al., 2012, 2014; Vandermosten et al., 2019). It further suggests that DD might be due to early abnormality in the language network.

Furthermore, our conjunction analysis of the two language subgroups confirmed that the left STG is a site for a common deficit in both language groups with reduced GMV and decreased brain activation. The evidence suggests that this is a neural signature of DD. The STG is where the auditory cortex is located, which is responsible for auditory perception and phonological analysis of words and sentences (Friederici, 2012). Recently, Skeide et al. (2018) found hypermyelination in the left auditory cortex in readers with DD using ultra-high-field MRI at 7T, and disrupted neural firing induced by hypermyelination in the layer IV of the auditory cortex, which may cause hypoactivation in the left STG. In summary, we found structural and functional alterations at the left STG in individuals with DD regardless of languages, which supports the phonological deficit hypothesis.

However, we failed to find consistent structural and functional deficits in the OT area. The main reason was that there was no structural alteration but only functional reduction at this area. The OT area is a key region for orthographic recognition during visual word processing (Glezer et al., 2009, 2016; Hirshorn et al., 2016; Nobre et al., 1994) and was reported to be impaired in individuals with DD (McCrory et al., 2005; Richlan et al., 2010; Wandell et al., 2012). The specialization of this region for orthographic processing is developed along with reading acquisition (Brem et al., 2010), and the dysfunction of the OT area in DD is possibly a result of reading failure (Pugh et al., 2000). A recent meta-analysis of VBM studies (McGrath & Stoodley, 2019) also failed to detect structural deficit in the OT area, which is consistent with our finding. Taken together, the lack of structural deficits with only hypoactivation at the OT area appears to suggest that the visuo-orthographic deficits at the OT might be a consequence of being DD. In contrast, the left STG which was discussed above, appears to be associated with the cause of DD. Our results provide further support for the phonological deficit hypothesis that phonological deficit is the primary deficit and other deficits may be a result of the phonological deficit (Pugh et al., 2000).

### Language differences in structural and functional alterations

The left dorsal IFG which peaked at (-48, 8, 26) showed greater reduction in morpho-syllabic languages than in alphabetic languages for both GMV and brain activation, suggesting greater impairment in this region in morpho-syllabic languages than in alphabetic languages, and the greater activation reduction in morpho-syllabic languages was also verified in the confirmation analysis. Previously, many Chinese studies have reported impairment at the dorsal left IFG, for example, reduced brain activation in an auditory rhyming judgment task in children with DD at (-44, 10, 26) (Cao et al., 2017), in a lexical decision task at (-44, 3, 29) (Siok et al., 2004), in a homophone judgment task at (-55, 5, 22) (Siok et al., 2008), and in a morphological task at (-36, 8, 26) (Liu et al., 2013b). This left dorsal IFG has been believed to be more involved in Chinese reading than in alphabetic languages, with a peak at the left MFG (-46, 18, 28) as reported in a previous meta-analysis study (Tan et al., 2005). The dorsal IFG was found to be involved in phonological processing in Chinese reading (Wu et al., 2012), and it is thought to be related to addressed phonology during Chinese character reading (Tan et al., 2005). Our study adds to the literature that not only the brain activation but also the GMV is more reduced in the left dorsal IFG in individuals with DD in morpho-syllabic languages than in alphabetic languages. This might be due to the fact that healthy Chinese readers have increased GMV and brain activation in the left dorsal IFG than healthy alphabetic readers, because the features of Chinese require greater involvement of this region in reading than alphabetic languages due to the whole-character-to-whole-syllable mapping. Actually, two cross-linguistic studies have argued that different findings of DD in different languages are actually driven by the fact that control readers show language-specific brain activation patterns (Feng et al., 2020; Hu et al., 2010), and that brain activation in individuals with DD is actually the same across languages. For example, Hu et al. (2010) found that Chinese control readers showed greater activation in the left IFG, and English control readers showed greater activation in the left superior temporal sulcus; however, children with DD in Chinese and English showed similar brain activation in these two regions. Therefore, readers with DD fail to show language specialization due to their limited reading experience and skills. In summary, this language-specific deficit is believed to be a consequence of being DD in learning morpho-syllabic languages, indicating their inability to accommodate to their own writing system.

In the direct comparison between alphabetic and morpho-syllabic languages, we also found greater hypoactivation in DD in alphabetic languages than in morpho-syllabic languages in the left MTG, right STG and left fusiform gyrus, which was verified in the confirmation analysis. Our finding is consistent with previous neuroimaging studies that revealed reduced activation associated with DD in the posterior reading network in alphabetic languages (Paulesu et al., 2001; Richlan et al., 2010; Vandermosten et al., 2019), suggesting deficient orthographic and phonological processing. However, the novelty of the current study is to demonstrate greater severity of deficit in these regions in alphabetic languages than in morpho-syllabic languages. In a previous meta-analysis study of alphabetic languages, it was found that there was greater hypoactivation in the left fusiform gyrus (-40, -42, -16) in shallow orthographies than in deep orthographies (Martin et al., 2016). The explanation is that this region is associated with bottom-up rapid processing of letters, because it was found that at a proximal region (-38, -50, -16), there was a word length effect for German nonwords in non-impaired readers (Schurz et al., 2010). Moreover, the left fusiform gyrus has been found to be more involved in English reading than in Chinese reading in typical mature readers with a peak of the effect at (-44, -56, -12) (Tan et al., 2005). Therefore, the left fusiform gyrus is important for letter-by-letter orthographic recognition in alphabetic languages, and this explains why we found greater deficit in this region in alphabetic languages than in morpho-syllabic languages. As for the right STG, the previous meta-analysis found greater hypoactivation in deep orthographies than shallow orthographies (Martin et al., 2016). Together with our finding, it suggests that the right STG might be associated with the inconsistent mapping between graphemes and phonemes in deep orthographic alphabetic languages. In summary, these greater deficits in alphabetic languages than in morpho-syllabic languages might be due to the inability to adapt to the special features of alphabetic languages in individuals with DD.

For the structural studies, we found greater GMV alterations in morpho-syllabic languages than in alphabetic languages, including greater GMV reduction in the left STG, left MFG, left supramarginal gyrus, left SOG and left insula, as well as greater GMV increase in the right STG and ITG. However, considering the limited number of studies included in the morpho-syllabic language group and inconsistent results with functional studies, the results should be interpreted with caution.

### Compensation mechanism

In the multi-modal meta-analysis, we found increased GMV and hyperactivation in participants with DD in the right MTG which was mainly driven by the morpho-syllabic languages. For the functional studies, we also found greater hyperactivation in the right precentral gyrus in morpho-syllabic languages than in alphabetic languages. We believe that these findings are related to the compensation mechanism of the right hemisphere. As precentral gyri play an important role in articulation (Dronkers, 1996), overactivation in the precentral gyrus is interpreted as an articulation strategy used by individuals with DD to compensate for their deficient phonological processing (Cao et al., 2018; Shaywitz et al., 1998; Waldie et al., 2013). The compensation in the right MTG is developed in morpho-syllabic languages presumably due to the tight connection between orthography and semantics (Wang et al., 2015). Substantial evidence has shown that dyslexia was often accompanied by excessive activation of the right hemisphere (Cao et al., 2017, 2018; Kovelman et al., 2012; Kronschnabel et al., 2014; Yang & Tan, 2020) and reduced left lateralization of the language network (Altarelli et al., 2014; Bloom et al., 2013). Furthermore, training studies have also found increased activation in many regions in the right hemisphere in individuals with DD after reading intervention (Barquero et al., 2014; Meyler et al., 2008), suggesting the compensatory role of the right hemisphere when the left language/reading network is deficient (Coslett & Monsul, 1994; Weiller et al., 1995). However, according to the previous meta-analysis study, different regions showed overactivation in different writing systems (Martin et al., 2016). In particular, the left anterior insula showed greater overactivation in deep orthographies while the left precentral gyrus showed greater overactivation in shallow orthographies in individuals with DD. Taken together, it suggests that different compensatory mechanisms are developed depending on the characteristics of the writing system as well as learning experiences, and the compensation in the right MTG and right precentral gyrus appears to be particularly salient in morpho-syllabic languages.

### Divergent structural and functional alterations in DD

In the multimodal analysis, we also found divergent structural and functional changes related to DD, including the left IPL and left cerebellum where there was increased GMV and hypoactivation and bilateral caudate where there was reduced GMV and hyperactivation. These divergent structural and functional alterations in DD were primarily driven by alphabetic languages.

### The left IPL

We found increased GMV and hypoactivation in DD in the left IPL. Consistent hypoactivation in the left IPL in DD has been documented in previous studies (Maisog et al., 2008; Martin et al., 2016). Furthermore, it was found that the deficit of the left IPL was greater in children than in adults with DD (Richlan et al., 2011), suggesting that the functional impairment of the left IPL may gradually recover with development. This may be related to the transfer of the reading circuit from the dorsal pathway to the ventral pathway over development (Younger et al., 2017). Control children activate the left IPL to a greater degree than control adults because they rely more on the dorsal pathway. Therefore, children with DD show a great reduction in the left IPL in comparison to adults with DD. Alternatively, the left IPL has been found to be deactivated during language tasks (Cao et al., 2008, 2017; Meyler et al., 2008; Schulz et al., 2009), and this is due to the nature of the default mode network (Laird et al., 2009), which is deactivated during active tasks. Therefore, it might be the case that the increased GMV in individuals with DD increases inhibitory inputs received by the IPL, which results in greater deactivation.

For structural studies involving the left IPL, the results are inconsistent. The GMV of the left supramarginal gyrus around the posterior part of perisylvian cortex was found to be reduced in individuals with DD (Linkersdorfer et al., 2012; McGrath & Stoodley, 2019) and it showed a positive correlation with reading accuracy only in normal readers (Jednorog et al., 2015). However, the GMV of the left inferior parietal cortex excluding the supramarginal and the angular was found to increase in individuals with DD (McGrath & Stoodley, 2019) and a study showed that the volume of the left inferior parietal cortex in control readers was negatively correlated with reading level (Houston et al., 2014). The IPL in the current study is outside the supramarginal and angular gyrus; therefore, it is consistent with the previous findings that there is increased GMV in individuals with DD.

### The cerebella

Increased GMV and hypoactivation in DD were also found in the left cerebellum; however, in the right cerebellum, for the alphabetic language group, we found decreased GMV and hyperactivation. Previously, it was found that the right cerebellum is greater in size than the left cerebellum in healthy controls while the asymmetry is reduced in individuals with DD (Kibby et al., 2008; Rae et al., 2002). This is consistent with our finding of increased GMV in the left cerebellum and decreased GMV in the right cerebellum in individuals with DD, suggesting reduced asymmetry in cerebellum.

The cerebella have been found to play an important role in inner speech, automatization in reading and suppression of overt articulatory movement in silent reading (Ait Khelifa-Gallois et al., 2015). Functional abnormality of cerebellum in DD has been reported repeatedly; however, hyperactivation was reported more often in the right cerebellum (Feng et al., 2017; Hernandez et al., 2013; Kronschnabel et al., 2014; Richlan et al., 2010; Rumsey et al., 1997; van Ermingen-Marbach et al., 2013a), while hypoactivation was reported more often in the left cerebellum (Christodoulou et al., 2014; McCrory et al., 2000; Olulade et al., 2012; Reilhac et al., 2013; Siok et al., 2008). This is consistent with our finding of hyperactivation in the right cerebellum and hypoactivation in the left cerebellum. Different alteration patterns in the left and right cerebellum suggest that they may play different roles in reading and dyslexia. The right cerebellum has been found to be connected with the left frontal-parietal pathway for phonological processing and with the left frontal-temporal pathway for semantic processing (Alvarez & Fiez, 2018; Gatti et al., 2020). The left cerebellum, however, is involved in error monitoring during reading unfamiliar non-words (Ben-Yehudah & Fiez, 2008), as well as articulation related movement process, since it is activated in reading aloud but not in lexical decision (Carreiras et al., 2007). Richards et al. (2006) argued that the left cerebellum is involved in processing the morphology of word forms, and the right cerebellum is involved in phonological processing. Therefore, hyperactivation in the right cerebellum in readers with DD suggests that they may use it as a compensation for their deficient phonological processing, while hypoactivation in the left cerebellum may suggest reduced error monitoring in readers with DD. Taken together, the finding of neurological alterations in the cerebellum supports the cerebellar deficit hypothesis (Stoodley & Stein, 2011), implicating the necessity of considering DD from a broader spectrum of developmental disorders.

It is still unclear why there is increased GMV but decreased activation in some brain regions. It may be due to the following reasons: (1) increased dendrites receiving more inhibitory input from other neurons; (2) abnormal neuronal migration deactivated the firing of neurons as a result of disrupted local microcircuits (Giraud & Ramus, 2013); (3) weaker input from other regions deactivated the target region and changed the structure of the region (Wang et al., 2019).

### The caudate

We also found decreased GMV and hyperactivation in readers with DD in the bilateral caudate. Previous studies have reported decreased GMV (Brown et al., 2001; Jagger-Rickels et al., 2018; McGrath & Stoodley, 2019; Tamboer et al., 2015) and hyperactivation in bilateral caudate in individuals with DD (Martin et al., 2016; Olulade et al., 2012; Pekkola et al., 2006; Richlan et al., 2010, 2011; Rumsey et al., 1997), which is consistent with our finding. Furthermore, the GMV volume of the caudate in individuals with DD was found to be positively correlated with reading performance (Pernet et al., 2009; Tamboer et al., 2015), and the left caudate’s activation was correlated with longer reaction time in word reading only in individuals with DD (Cheema et al., 2018). The caudate plays an important role in procedural learning and phonological processing (Grahn et al., 2008; Tettamanti et al., 2005; Ullman et al., 2020). Decreased GMV and increased activation at the bilateral caudate might be caused by reduced dendrites and reduced inhibitory inputs received in individuals with DD (Achal et al., 2016; Finn et al., 2014). It may also be due to pre-existing local structural deficit leading to compensatory hyperactivation of the remaining part of the caudate. GMV reduction in basal ganglia was found in many other neuropsychiatric disorders, such as attention-deficit hyperactivity disorder (Frodl & Skokauskas, 2012; Mous et al., 2015; Nakao et al., 2011), autism spectrum disorder (Nickl-Jockschat et al., 2012) and major depression disorder (Husain et al., 1991; Lu et al., 2016). Altered myelination and neurotransmitters may contribute to the structural and functional alterations related to basal ganglia (Nord et al., 2019; Wichmann & DeLong, 2012).

In addition, there are also brain regions that only showed alteration in structure but not in function or vice versa. It suggests that brain activation is only partially determined by GMV and that there are other factors influencing brain activation which might include but are not limited to myelination, synaptic processes, as well as interconnectivities between regions.

## Limitation

In this meta-analysis, we found convergent and divergent functional and structural alterations across writing systems. However, due to limitations of neuroimaging techniques and the lack of evidence from multi-scale studies, the neurophysiology of changes in brain structure and function is unknown.

## Conclusion

We found convergent functional and structural alterations in the left STG across different writing systems, suggesting a neural signature of DD, which might be associated with phonological deficit. We also found greater functional and structural alteration in the left dorsal IFG in morpho-syllabic languages than alphabetic languages, suggesting a language-specific effect of DD, which might be related to the special feature of whole-character-to-whole-syllable mapping in morpho-syllabic languages.

## Acknowledgments

This work was supported by the “Fundamental Research Funds for the Central Universities” awarded to Dr. Fan Cao, “Guangdong Planning Office of Philosophy and Social Science” (GD19CXL05) awarded to Dr. Fan Cao, and by “Science and Technology Program of Guangzhou, China, Key Area Research and Development Program (202007030011)”.

---

**Table S1.** Convergent functional deficits in individuals with DD across all languages (JK represents the results of jack-knife sensitivity analysis)

| Regions | MNI coordinate | SDM-Z | P | Voxels | Cluster breakdown (Voxels) | JK |
| --- | --- | --- | --- | --- | --- | --- |
| *Hypoactivation in DD* |  |  |  |  |  |  |
| Left IPL | -54,-44,30 | 5.427 | 0.0000 | 12510 | Left IPL, BA 40 (1011)<br>Left MTG, BA 21 (694)<br>Left MTG, BA 37 (620)<br>Left ITG, BA 37 (608)<br>Left fusiform gyrus, BA 37 (467)<br>Left inferior network, inferior longitudinal fasciculus (429)<br>Left ITG, BA 20 (389)<br>Left STG, BA 48 (372)<br>Left supramarginal gyrus, BA 48 (354)<br>Left rolandic operculum, BA 48 (314)<br>Left angular gyrus, BA 39 (312)<br>Left MTG, BA 22 (301)<br>Left STG, BA 42 (290)<br>Left precentral gyrus, BA 6 (271)<br>Left arcuate network, posterior segment (261)<br>Left cerebellum, lobule VI, BA 37 (260)<br>Left IFG, opercular part, BA 44 (256)<br>Left superior parietal gyrus, BA 7 (250)<br>Left superior longitudinal fasciculus III (229)<br>Left STG, BA 22 (215)<br>Left cerebellum, crus I, BA 37 (214)<br>Left supramarginal gyrus, BA 40 (202)<br>Left IFG, triangular part, BA 45 (197)<br>Left IPL, BA 2 (194)<br>Left fusiform gyrus, BA 19 (171) | 92/92 |
|  |  |  |  |  | Left IFG, opercular part, BA 48 (170)<br>Corpus callosum (159)<br>Left precentral gyrus, BA 44 (155) |  |
| Right middle occipital gyrus | 42,-86,6 | 1.988 | 0.0008 | 178 |  | 91/92 |
| *Hyperactivation in DD* |  |  |  |  |  |  |
| Right cerebellum | 24,-56,-30 | -1.500 | 0.0000 | 1968 | Right cerebellum, lobule VI, BA 37 (359)<br>Middle cerebellar peduncles (313)<br>Right cerebellum, lobule VI, BA 19 (172)<br>Right cerebellum, lobule VI (153) | 92/92 |
| Right postcentral gyrus | 40,-26,56 | -1.347 | 0.0001 | 951 | Right postcentral gyrus, BA 3 (246)<br>Right precentral gyrus, BA 4 (238) | 92/92 |
| Right caudate nucleus | 10,2,14 | -1.224 | 0.0002 | 450 | Right anterior thalamic projections (176)<br>Right caudate nucleus (157) | 92/92 |
| Left caudate nucleus | -16,12,6 | -1.253 | 0.0002 | 413 | Left anterior thalamic projections (269) | 92/92 |

**Table S2.** Convergent structural deficits in individuals with DD across all languages (JK represents the results of jack-knife sensitivity analysis)

| Regions | MNI coordinate | SDM-Z | P | Voxels | Cluster breakdown (Voxels) | JK |
| --- | --- | --- | --- | --- | --- | --- |
| *Decreased GMV in DD* |  |  |  |  |  |  |
| Left IFG | -38,44,-16 | 2.275 | 0.0001 | 494 | Left IFG, orbital part, BA 47 (182) | 26/27 |
| Right STG | 56,-44,18 | 2.027 | 0.0006 | 365 |  | 25/27 |
| *Increased GMV in DD* |  |  |  |  |  |  |
| Right MTG | 46,-4,-22 | -1.658 | 0.0001 | 747 | Right inferior network (210) | 26/27 |
| Left IPL | -42,-36,36 | -1.817 | 0.0000 | 235 | Left IPL, BA 40 (151) | 26/27 |

**Table S3.**
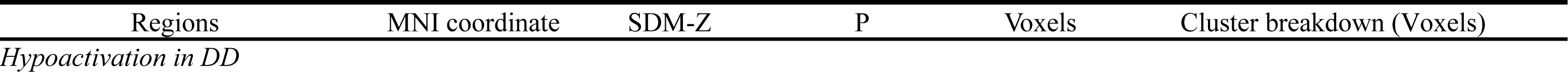
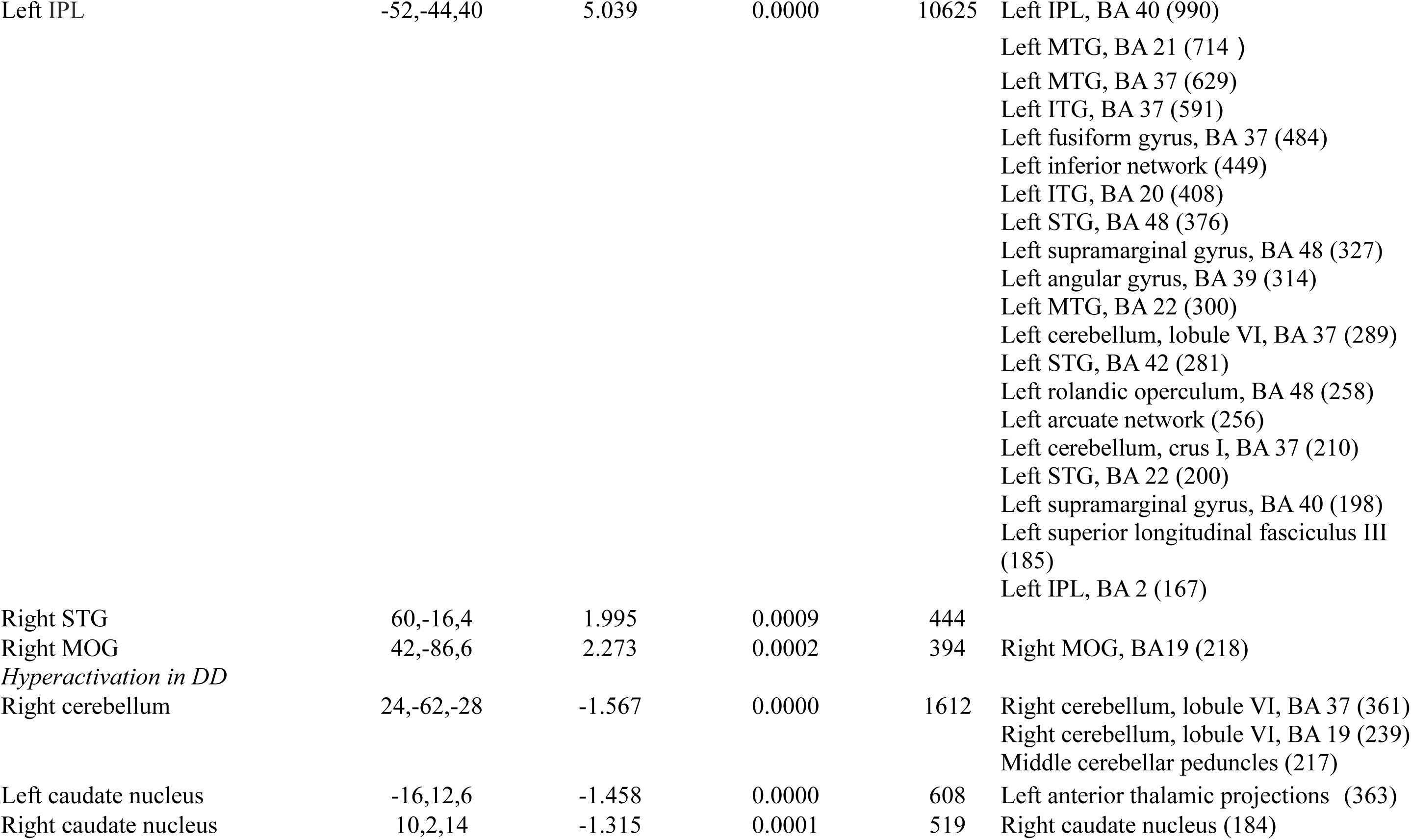
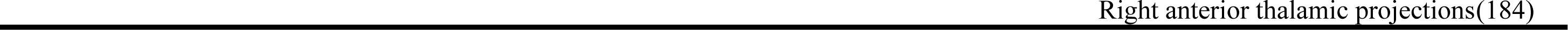
Convergent functional deficits in individuals with DD across alphabetic languages.

| Regions | MNI coordinate | SDM-Z | P | Voxels | Cluster breakdown (Voxels) |
| --- | --- | --- | --- | --- | --- |
| *Hypoactivation in DD* |  |  |  |  |  |
| Left IPL | -52,-44,40 | 5.039 | 0.0000 | 10625 | Left IPL, BA 40 (990)<br>Left MTG, BA 21 (714 )<br>Left MTG, BA 37 (629)<br>Left ITG, BA 37 (591)<br>Left fusiform gyrus, BA 37 (484)<br>Left inferior network (449)<br>Left ITG, BA 20 (408)<br>Left STG, BA 48 (376)<br>Left supramarginal gyrus, BA 48 (327)<br>Left angular gyrus, BA 39 (314)<br>Left MTG, BA 22 (300)<br>Left cerebellum, lobule VI, BA 37 (289)<br>Left STG, BA 42 (281)<br>Left rolandic operculum, BA 48 (258)<br>Left arcuate network (256)<br>Left cerebellum, crus I, BA 37 (210)<br>Left STG, BA 22 (200)<br>Left supramarginal gyrus, BA 40 (198)<br>Left superior longitudinal fasciculus III (185)<br>Left IPL, BA 2 (167) |
| Right STG | 60,-16,4 | 1.995 | 0.0009 | 444 |  |
| Right MOG | 42,-86,6 | 2.273 | 0.0002 | 394 | Right MOG, BA19 (218) |
| *Hyperactivation in DD* |  |  |  |  |  |
| Right cerebellum | 24,-62,-28 | -1.567 | 0.0000 | 1612 | Right cerebellum, lobule VI, BA 37 (361)<br>Right cerebellum, lobule VI, BA 19 (239)<br>Middle cerebellar peduncles (217) |
| Left caudate nucleus | -16,12,6 | -1.458 | 0.0000 | 608 | Left anterior thalamic projections (363) |
| Right caudate nucleus | 10,2,14 | -1.315 | 0.0001 | 519 | Right caudate nucleus (184) |

**Table S4.** Convergent structural deficits in individuals with DD across alphabetic languages.

| Regions | MNI coordinate | SDM-Z | P | Voxels | Cluster breakdown (Voxels) |
| --- | --- | --- | --- | --- | --- |
| *Decreased GMV in DD* |  |  |  |  |  |
| Left IFG | -38,42,-16 | 2.306 | 0.0001 | 611 | Left IFG, orbital part, BA 47 (217) |
| Right STG | 56,-44,18 | 2.024 | 0.0003 | 560 | Right STG, BA 42 (156) |
| Right caudate | 6,14,2 | 1.695 | 0.0022 | 166 |  |
| *Increased GMV in DD* |  |  |  |  |  |
| Left IPL | -42,-36,36 | -1.976 | 0.0000 | 237 | Left IPL, BA 40 (237) |
| Right MTG | 50,-12,-14 | -1.040 | 0.0014 | 174 |  |

**Table S5.**
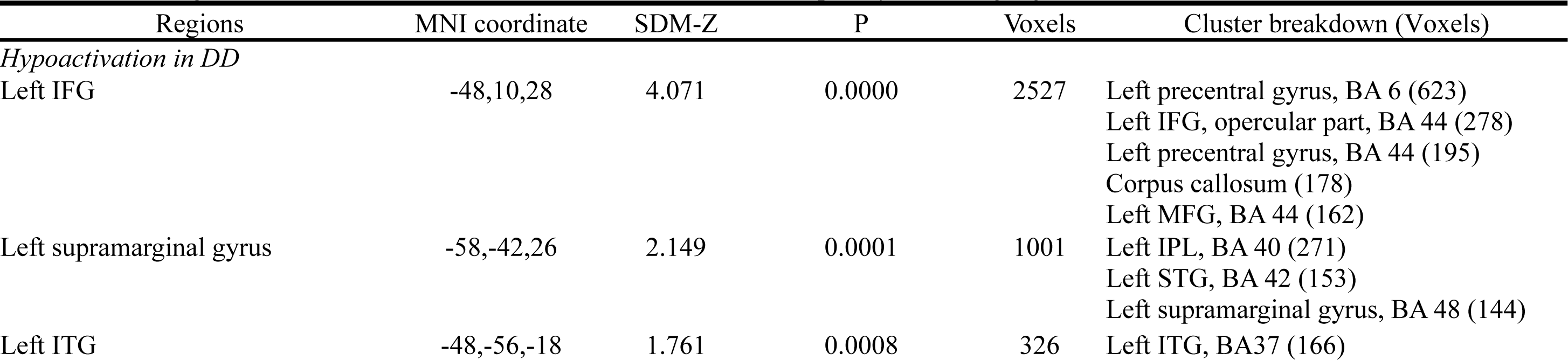
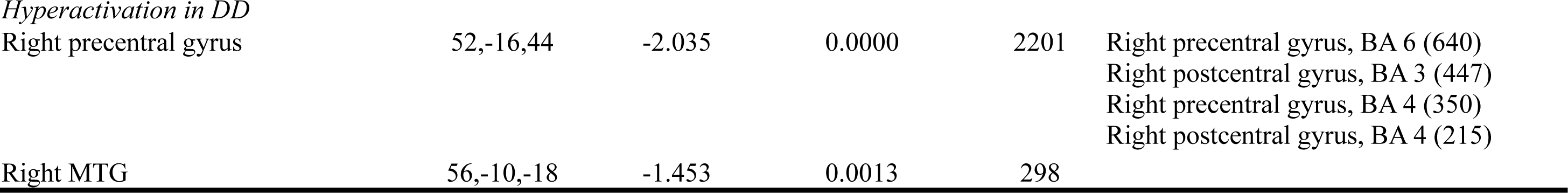
Convergent functional deficits in individuals with DD across morpho-syllabic languages.

| Regions | MNI coordinate | SDM-Z | P | Voxels | Cluster breakdown (Voxels) |
| --- | --- | --- | --- | --- | --- |
| *Hypoactivation in DD* |  |  |  |  |  |
| Left IFG | -48,10,28 | 4.071 | 0.0000 | 2527 | Left precentral gyrus, BA 6 (623)<br>Left IFG, opercular part, BA 44 (278)<br>Left precentral gyrus, BA 44 (195)<br>Corpus callosum (178)<br>Left MFG, BA 44 (162) |
| Left supramarginal gyrus | -58,-42,26 | 2.149 | 0.0001 | 1001 | Left IPL, BA 40 (271)<br>Left STG, BA 42 (153)<br>Left supramarginal gyrus, BA 48 (144) |
| Left ITG | -48,-56,-18 | 1.761 | 0.0008 | 326 | Left ITG, BA37 (166) |
| *Hyperactivation in DD* |  |  |  |  |  |
| Right precentral gyrus | 52,-16,44 | -2.035 | 0.0000 | 2201 | Right precentral gyrus, BA 6 (640)<br>Right postcentral gyrus, BA 3 (447)<br>Right precentral gyrus, BA 4 (350)<br>Right postcentral gyrus, BA 4 (215) |
| Right MTG | 56,-10,-18 | -1.453 | 0.0013 | 298 |  |

**Table S6.** Convergent structural deficits in individuals with DD across morpho-syllabic languages.

| Regions | MNI coordinate | SDM-Z | P | Voxels | Cluster breakdown (Voxels) |
| --- | --- | --- | --- | --- | --- |
| *Decreased GMV in DD* |  |  |  |  |  |
| Left STG | -50,4,-4 | 2.466 | 0.0000 | 2948 | Left insula, BA 48 (539)<br>Left STG, BA 38 (392)<br>Left rolandic operculum, BA 48 (226)<br>Left MTG, BA 21 (215)<br>Left STG, BA 48 (186) |
| Left temporoparietal cortex | -56,-40,18 | 2.102 | 0.0002 | 900 | Left supramarginal gyrus, BA 48 (188)<br>Left STG, BA 42 (171) |
| Left Calcarine Cortex | -20,-66,14 | 2.447 | 0.0000 | 449 | Corpus callosum (297) |
| Left MFG | -32,26,40 | 2.319 | 0.0001 | 438 |  |
| *Increased GMV in DD* |  |  |  |  |  |
| Right STG | 34,6,-26 | -1.572 | 0.0001 | 1829 | Right inferior network (273)<br>Right STG, BA 38 (261)<br>Right ITG, BA 20 (250)<br>Right MTG, BA 20 (212) |
| Right precuneus | 12,-52,42 | -1.254 | 0.0014 | 156 |  |

**Table S7.** Convergent deficits in individuals with DD only in functional studies not in structural studies.

| Regions | MNI coordinate | SDM-Z | Voxels | Cluster breakdown (Voxels) |
| --- | --- | --- | --- | --- |
| *Hypoactivation in DD* |  |  |  |  |
| Left supramarginal gyrus | -56,-44,38 | 5.581 | 7946 | Left MTG (1694)<br>Left ITG (1108)<br>Left fusiform gyrus (745)<br>Left IPL (496)<br>Left supramarginal gyrus (428)<br>Left STG (372)<br>Left precentral gyrus (366)<br>Left inferior occipital gyrus (312)<br>Left angular gyrus (282)<br>Left superior Parietal gyrus (275)<br>Left IFG (274)<br>Left cerebellum Crus (256)<br>Left IFG (213) |
| Right middle occipital gyrus | 42,-86,6 | 1.988 | 178 |  |
| *Hyperactivation in DD* |  |  |  |  |
| Right Cerebellum | 24,-56,-30 | -1.500 | 1968 | Right cerebellum, Anterior Lobe (1356) |
| Right precentral gyrus | 46,-26,56 | -1.347 | 951 | Right postcentral gyrus (466)<br>Right precentral gyrus (437) |

**Table S8.** Convergent deficits in individuals with DD only in structural studies not in functional studies.

| Regions | MNI coordinate | SDM-Z | Voxels | Cluster breakdown (Voxels) |
| --- | --- | --- | --- | --- |
| *Decreased GMV in DD* |  |  |  |  |
| Left IFG | -38,44,-16 | 2.275 | 362 | Left IFG, orbital part (190) |
| Right STG | 56,-44,18 | 2.027 | 365 | Right STG, BA 42 (196) |
| *Increased GMV in DD* |  |  |  |  |
| Right MTG | 42,-2,-20 | -1.584 | 582 | Right MTG (154) |

**Table S9.**
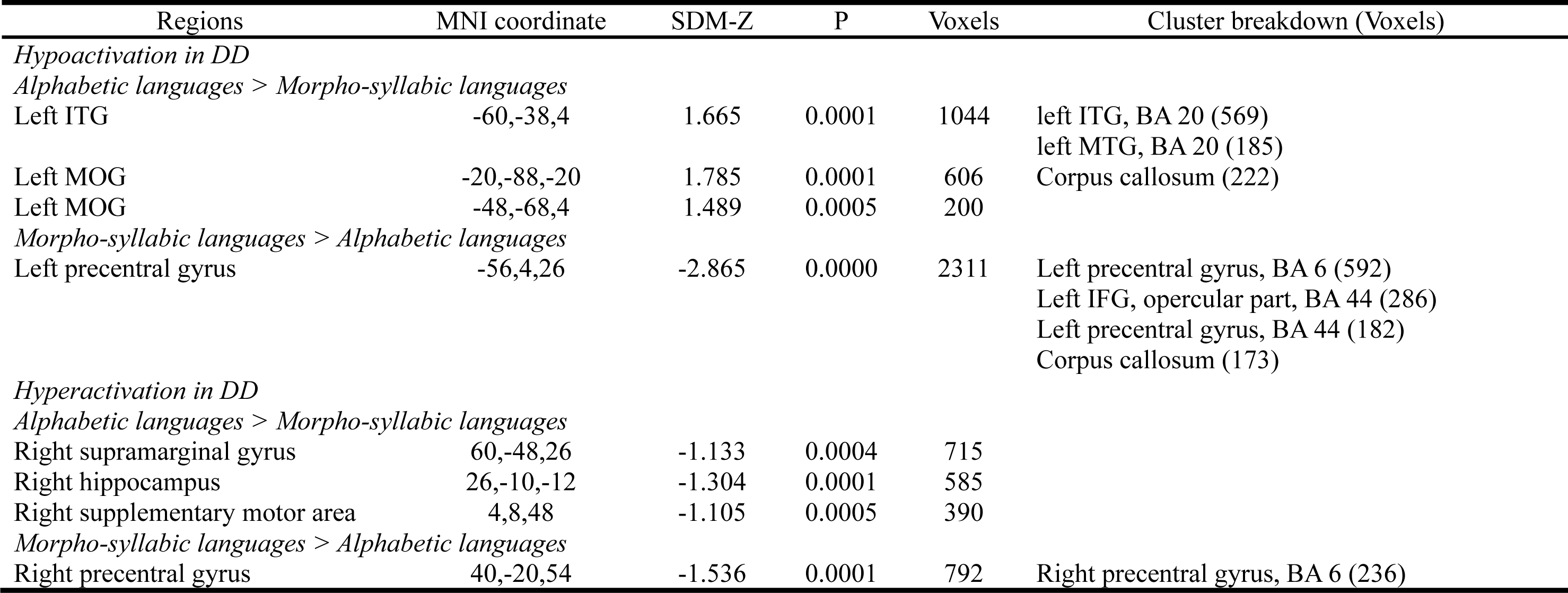
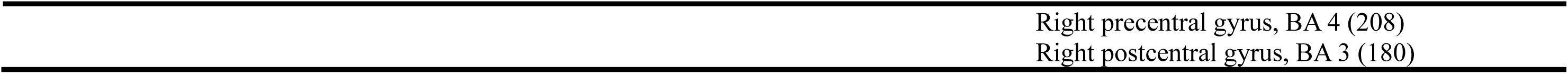
Confirmation study results when matched language, age and number of studies.

| Regions | MNI coordinate | SDM-Z | P | Voxels | Cluster breakdown (Voxels) |
| --- | --- | --- | --- | --- | --- |
| *Hypoactivation in DD* |  |  |  |  |  |
| *Alphabetic languages > Morpho-syllabic languages* |  |  |  |  |  |
| Left ITG | -60,-38,4 | 1.665 | 0.0001 | 1044 | left ITG, BA 20 (569)<br>left MTG, BA 20 (185) |
| Left MOG | -20,-88,-20 | 1.785 | 0.0001 | 606 | Corpus callosum (222) |
| Left MOG | -48,-68,4 | 1.489 | 0.0005 | 200 |  |
| *Morpho-syllabic languages > Alphabetic languages* |  |  |  |  |  |
| Left precentral gyrus | -56,4,26 | -2.865 | 0.0000 | 2311 | Left precentral gyrus, BA 6 (592)<br>Left IFG, opercular part, BA 44 (286)<br>Left precentral gyrus, BA 44 (182)<br>Corpus callosum (173) |
| *Hyperactivation in DD* |  |  |  |  |  |
| *Alphabetic languages > Morpho-syllabic languages* |  |  |  |  |  |
| Right supramarginal gyrus | 60,-48,26 | -1.133 | 0.0004 | 715 |  |
| Right hippocampus | 26,-10,-12 | -1.304 | 0.0001 | 585 |  |
| Right supplementary motor area | 4,8,48 | -1.105 | 0.0005 | 390 |  |
| *Morpho-syllabic languages > Alphabetic languages* |  |  |  |  |  |
| Right precentral gyrus | 40,-20,54 | -1.536 | 0.0001 | 792 | Right precentral gyrus, BA 6 (236) |

**Figure S1.**
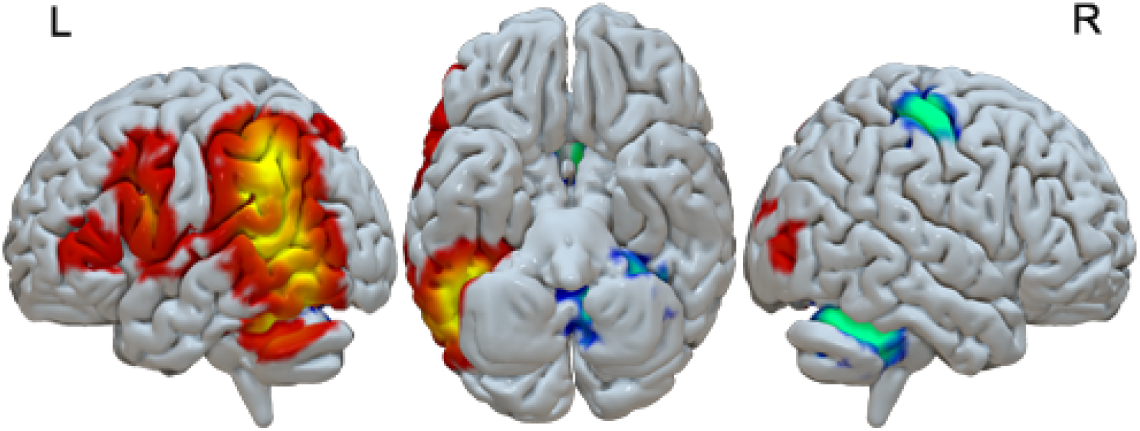
Functional deficits in individuals with DD across all languages (red-yellow: decreased activation in individuals with DD than in controls, blue-green: increased activation in individuals with DD than in controls).

**Figure S2.**
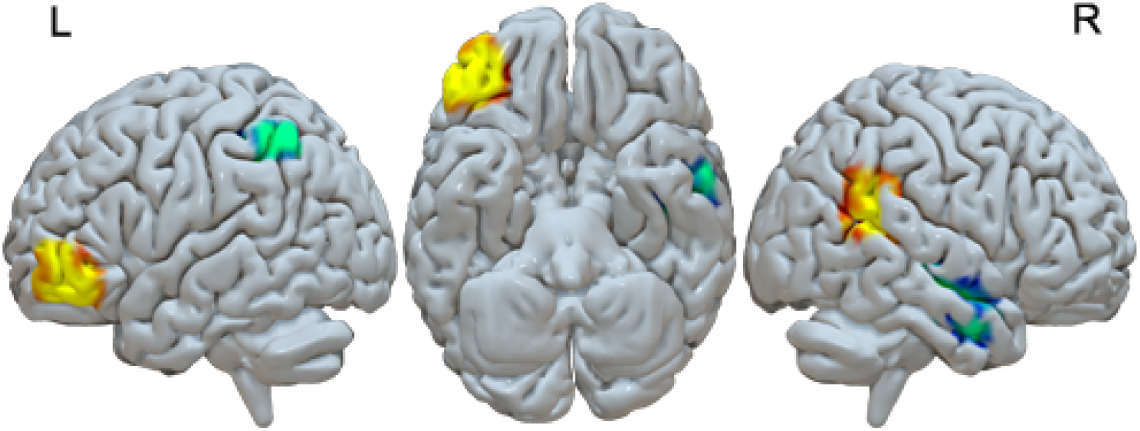
Structural deficits in individuals with DD across all languages (red-yellow: decreased GMV in individuals with DD than in controls, blue-green: increased GMV in individuals with DD than in controls).

**Figure S3.**
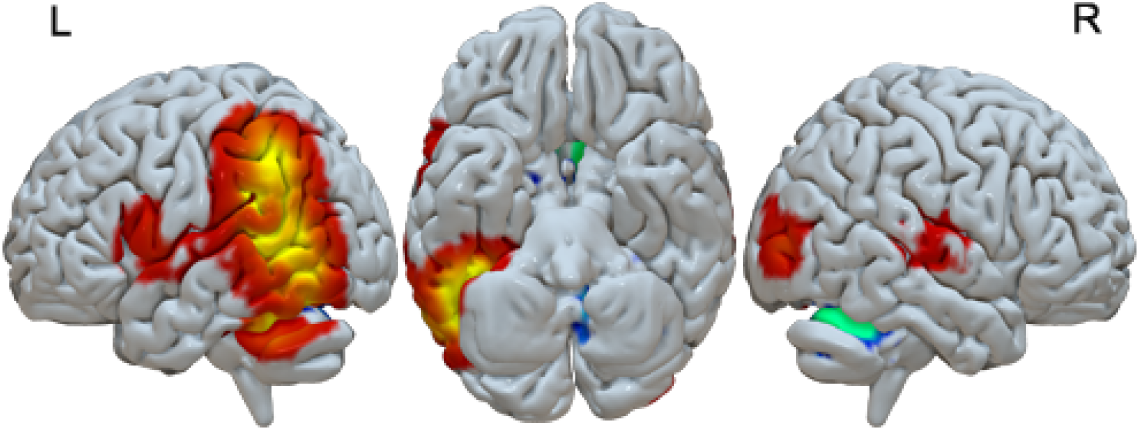
Functional deficits in individuals with DD in alphabetic languages (red-yellow: decreased activation in individuals with DD than in controls, blue-green: increased activation in individuals with DD than in controls).

**Figure S4.**
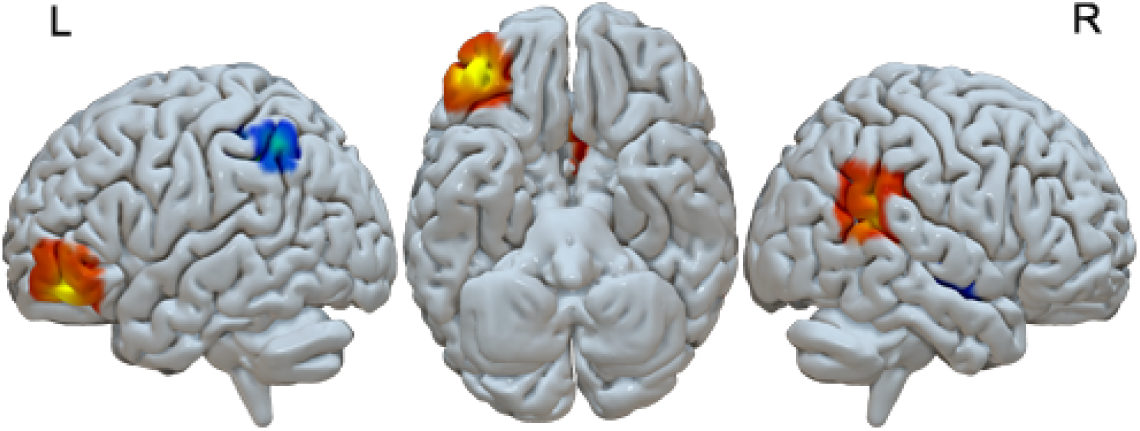
Structural deficits in individuals with DD in alphabetic languages (red-yellow: decreased GMV in individuals with DD than in controls, blue-green: increased GMV in individuals with DD than in controls).

**Figure S5.**
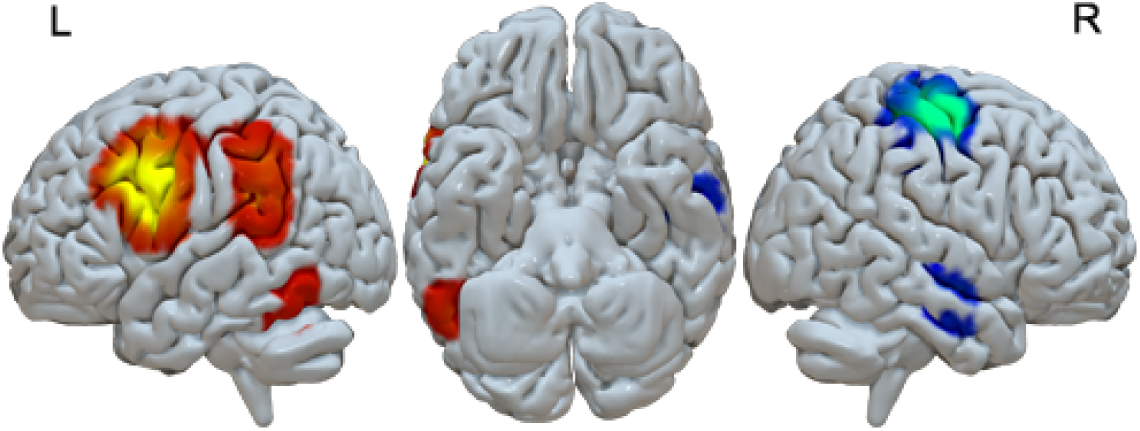
Functional deficits in individuals with DD in morpho-syllabic languages (red-yellow: decreased activation in individuals with DD than in controls, blue-green: increased activation in individuals with DD than in controls).

**Figure S6.**
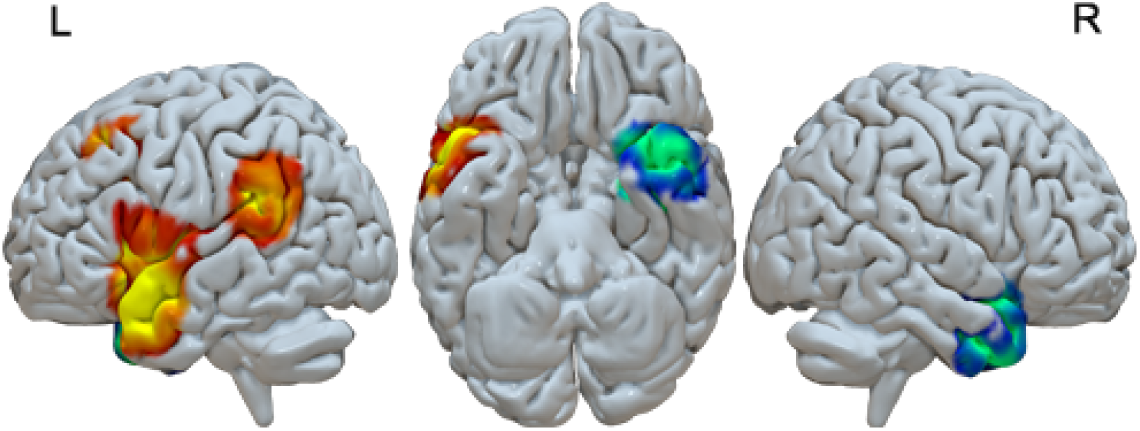
Structural deficits in individuals with DD in morpho-syllabic languages (red-yellow: decreased GMV in individuals with DD than in controls, blue-green: increased GMV in individuals with DD than in controls).

**Figure S7.**
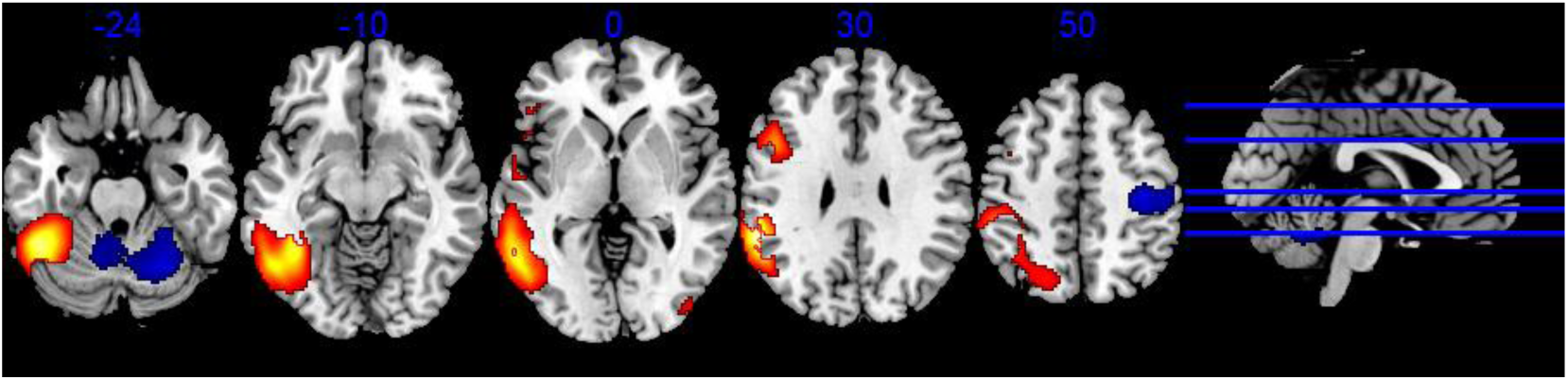
Convergent deficits in individuals with DD only in functional studies not in structural studies (red: TD>DD, blue: TD<DD).

**Figure S8.**
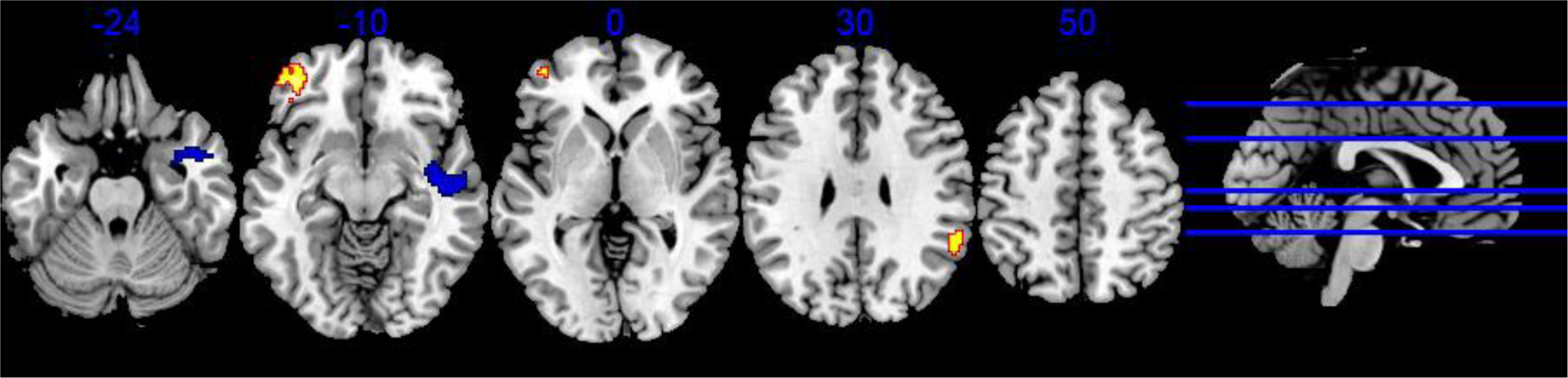
Convergent deficits in individuals with DD only in structural studies not in functional studies (red: TD>DD, blue: TD<DD).

**Figure S9.**
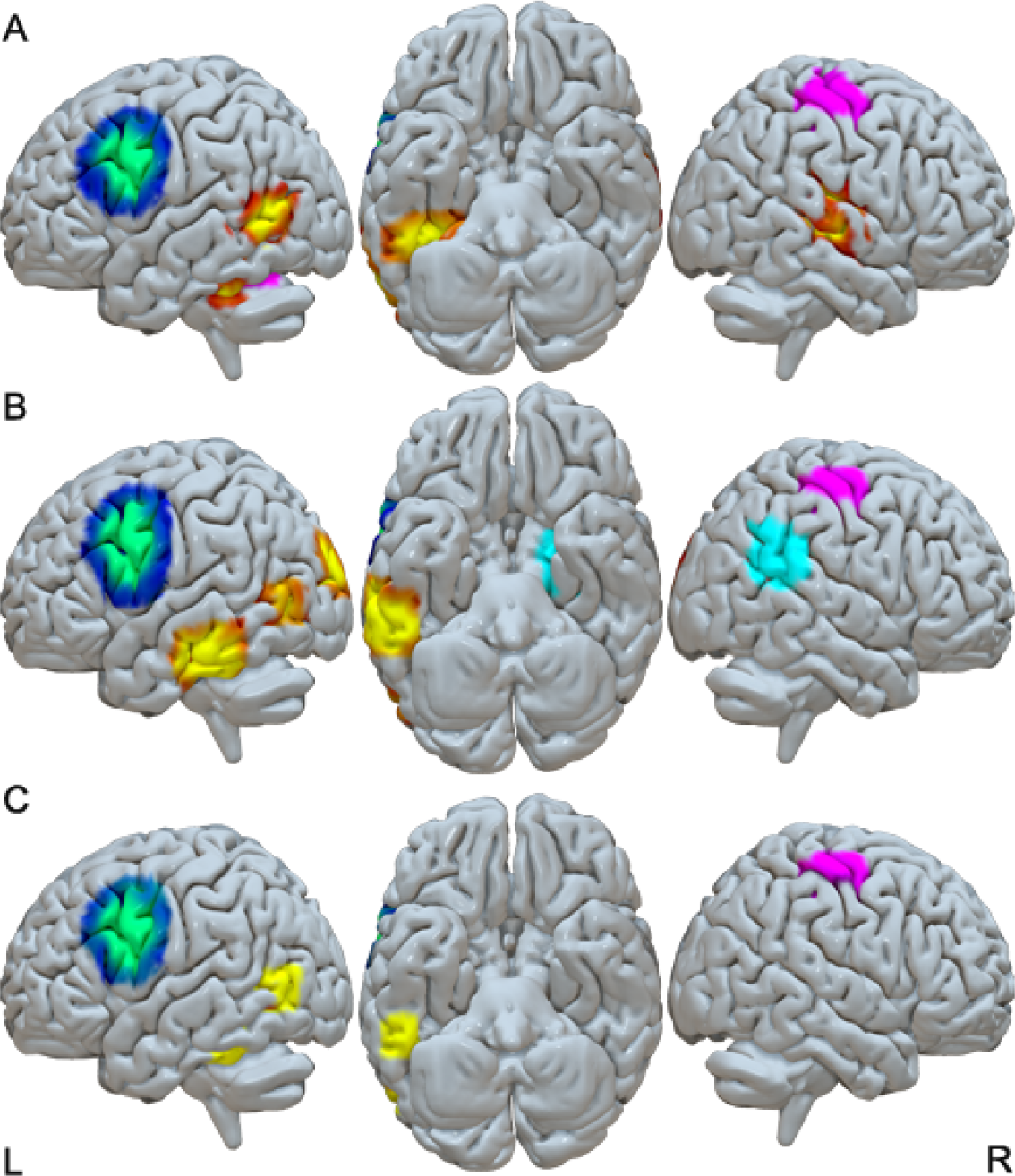
Direct comparison between alphabetic languages/English and morpho-syllabic languages/Chinese in fMRI/PET studies. Figure S9A. Direct comparison between alphabetic languages and morpho-syllabic languages. Figure S9B. Confirmation study results when matched language, age and number of studies. Figure S9C. Overlap of the two difference maps. (red-yellow: greater decreases in DD in alphabetic languages/English than in morpho-syllabic languages/Chinese; blue-green: greater decreases in DD in morpho-syllabic languages/Chinese than in alphabetic languages/English; purple: greater increases in DD in morpho-syllabic languages/Chinese than in alphabetic languages/English; cyan: greater increases in DD in alphabetic languages/English than in morpho-syllabic languages/Chinese).

## Notes

### Competing Interest Statement

The authors have declared no competing interest.

